# Integrated analysis of multimodal single-cell data

**DOI:** 10.1101/2020.10.12.335331

**Authors:** Yuhan Hao, Stephanie Hao, Erica Andersen-Nissen, William M. Mauck, Shiwei Zheng, Andrew Butler, Maddie J. Lee, Aaron J. Wilk, Charlotte Darby, Michael Zagar, Paul Hoffman, Marlon Stoeckius, Efthymia Papalexi, Eleni P. Mimitou, Jaison Jain, Avi Srivastava, Tim Stuart, Lamar B. Fleming, Bertrand Yeung, Angela J. Rogers, Juliana M. McElrath, Catherine A. Blish, Raphael Gottardo, Peter Smibert, Rahul Satija

## Abstract

The simultaneous measurement of multiple modalities, known as multimodal analysis, represents an exciting frontier for single-cell genomics and necessitates new computational methods that can define cellular states based on multiple data types. Here, we introduce ‘weighted-nearest neighbor’ analysis, an unsupervised framework to learn the relative utility of each data type in each cell, enabling an integrative analysis of multiple modalities. We apply our procedure to a CITE-seq dataset of hundreds of thousands of human white blood cells alongside a panel of 228 antibodies to construct a multimodal reference atlas of the circulating immune system. We demonstrate that integrative analysis substantially improves our ability to resolve cell states and validate the presence of previously unreported lymphoid subpopulations. Moreover, we demonstrate how to leverage this reference to rapidly map new datasets, and to interpret immune responses to vaccination and COVID-19. Our approach represents a broadly applicable strategy to analyze single-cell multimodal datasets, including paired measurements of RNA and chromatin state, and to look beyond the transcriptome towards a unified and multimodal definition of cellular identity.

**Availability:** Installation instructions, documentation, tutorials, and CITE-seq datasets are available at http://www.satijalab.org/seurat

## Introduction

The potential to catalog and characterize the rich diversity of cell types in the human immune system represents a powerful opportunity for single-cell genomics [1-5], yet also reveals the limitations of current approaches. While established technologies like single-cell RNA-seq (scRNA-seq) are capable of discovering new cell types and states in heterogeneous tissues, transcriptomics alone is often incapable of separating molecularly similar, but functionally distinct, categories of immune cells. Despite tremendous functional diversity, distinct populations of T cells such as effector, regulatory, γδ, and mucosal associated invariant T (MAIT), often cannot be effectively separated by scRNA-seq alone, even when using the most sensitive and cutting-edge technologies [6, 7]. This reflects technical challenges driven by the minimal RNA content of T cells coupled with high RNase expression [8-10], which hampers scRNA-seq data quality. More broadly, this exhibits the challenge of defining cell states based on the transcriptome alone, as important sources of cellular heterogeneity may not correlate strongly with transcriptomic features despite being identifiable in other modalities.

Multimodal single-cell technologies, which simultaneously profile multiple data types in the same cell, represent a new frontier for the discovery and characterization of cell states. For example, we recently introduced CITE-seq [11], which leverages oligonucleotide-conjugated antibodies to simultaneously quantify RNA and surface protein abundance in single cells via the sequencing of antibody-derived tags (ADTs). Moreover, pioneering technological advancements now enable the simultaneous profiling of transcriptome alongside either chromatin accessibility [12-14], DNA methylation [15, 16], nucleosome occupancy [17, 18], or spatial location [19, 20]. Each of these approaches offers an exciting solution to overcome the inherent limitations of scRNA-seq, and to explore how multiple cellular modalities affect cellular state and function [21].

The maturation of multimodal single-cell technologies also necessitates the development of new computational methods to integrate information across different data types [22]. For example, while CITE-seq datasets can be analyzed by first identifying clusters based on gene expression values [11, 23], and subsequently exploring their immunophenotypes, a multimodal computational workflow would define cell states based on both modalities. Importantly, these strategies must be robust to potentially large differences in the data quality and information content for each modality. In some contexts, robust protein quantifications may be most valuable for clustering, especially with a large and well-designed antibody panel. In other contexts (particularly when important cell type markers are missing or not previously known), the unsupervised nature of a cell’s transcriptome may be the most valuable. The varying information content of each modality, even across cells in the same dataset, represents a pressing challenge for the analysis and integration of multimodal datasets.

Here, we introduce ‘weighted-nearest neighbor’ (WNN) analysis, an analytical framework to integrate multiple data types measured within a cell, and to obtain a joint definition of cellular state. Our approach is based on an unsupervised strategy to learn cell-specific modality ‘weights’, which reflect the information content for each modality, and determine its relative importance in downstream analyses. We demonstrate that WNN analysis substantially improves our ability to define cellular states in multiple biological contexts and data types. We leverage this method to generate a multimodal ‘atlas’ based on a CITE-seq dataset of 211,000 human peripheral blood mononuclear cells (PBMC), with large cell-surface protein marker panels extending up to 228 antibodies. We utilize this dataset to identify and validate heterogeneous cell states in human lymphocytes and explore how the human immune system responds to vaccination and SARS-CoV-2 infection. Our approach, implemented in an updated version 4 of our open source R toolkit Seurat, represents a broadly applicable strategy for integrative multimodal analysis of single-cell data.

## Results

### Quantifying the relative utility of each modality in each cell

We sought to design a robust analytical workflow for the integration of multiple measurements collected within the same cell. To be applied to a range of biological contexts and data types, our strategy must successfully address the following criteria. First, the workflow must be robust to potentially vast differences in data quality between the modalities. Second, integrative multimodal analysis should enable multiple downstream analytical tasks, including visualization, clustering, and the identification of cellular trajectories. Lastly, and most importantly, simultaneous analysis of multiple modalities should improve on the ability to discover and characterize cell states, compared to independent analyses of each modality when performed separately.

These challenges highlight the importance of a flexible framework to handle diverse datasets. As previously described for CITE-seq [11, 24], the increased copy number of protein molecules compared to RNA molecules typically leads to more robust detection of protein features. The protein data in CITE-seq may therefore represent the most informative modality, particularly in cases where the antibody panel comprehensively represents all cell subsets with high specificity. Other panels may omit antibodies for key or previously undiscovered markers, or contain antibodies with low binding specificity, in which case the unsupervised nature of scRNA-seq may be most informative. Even within the same dataset, the relative utility of each modality to define cell states may vary across individual cells.

We therefore designed an analytical solution to address these goals, without requiring prior knowledge from the user regarding the importance of each modality. We first introduce and demonstrate our solution on our previously generated CITE-seq dataset of 8,617 cord blood mononuclear cells, with a panel of 10 immunophenotypic markers [11]. Independent unsupervised analysis of the RNA and protein data revealed largely consistent cell classifications (Figure 1A, B; Supplementary Figure 1), but did exhibit some differences. For example, CD8^+^ and CD4^+^ T cells were partially blended together when analyzing the transcriptome, but separated clearly in the protein data. Contrastingly, conventional dendritic cells (cDCs), along with a rare population of erythroid progenitors and spiked-in murine 3T3 controls, formed distinct clusters when analyzing RNA, but were intermixed with other cell types based on surface protein abundance. With biological foresight, the cell-type specific differences across modalities could be predicted by the composition of the CITE-seq panel, which contained anti-CD4 and anti-CD8 antibodies, but lacked any immunophenotypic markers to discriminate cDCs.

**Figure 1:**
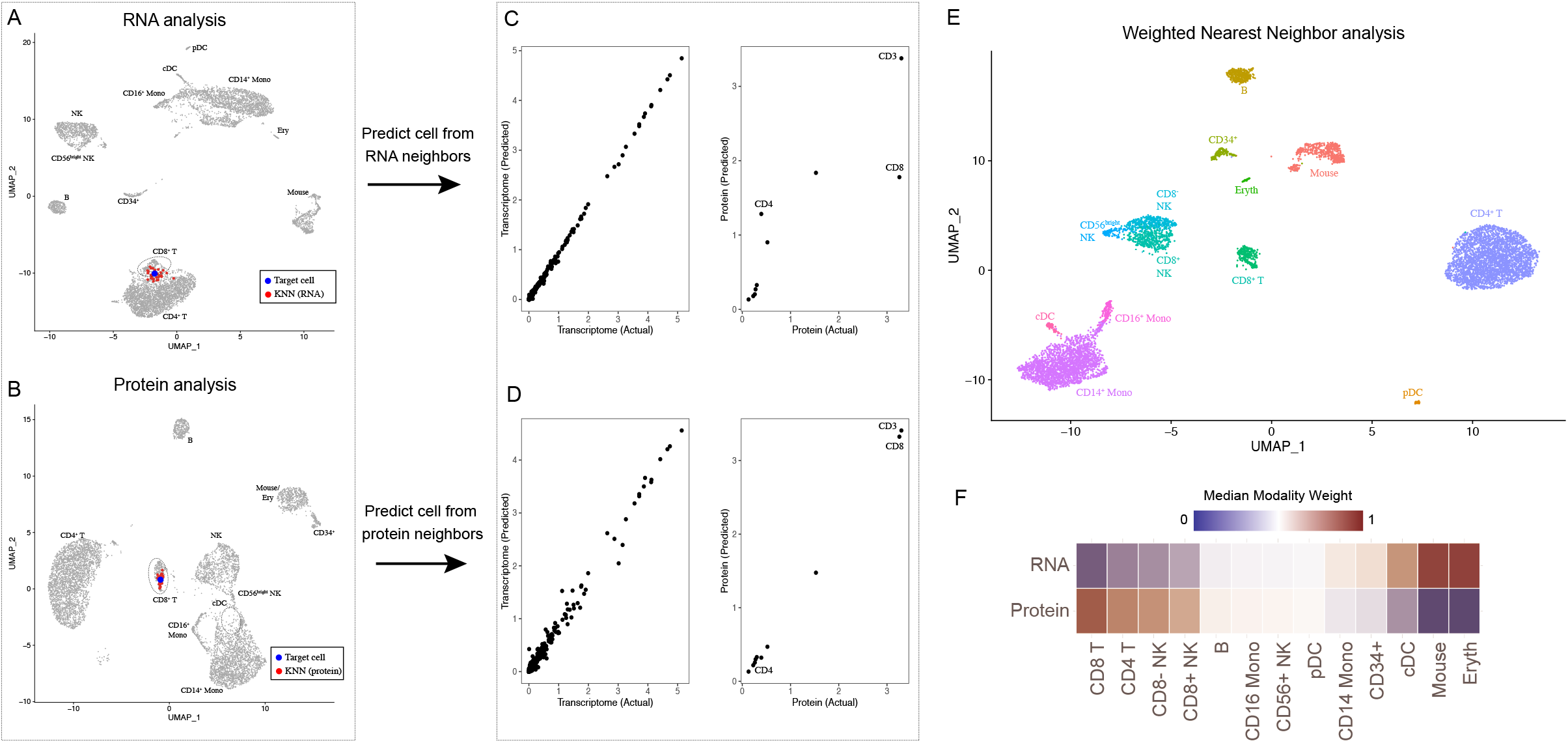
Schematic overview of multimodal integration using Weighted Nearest Neighbor analysis. **(A, B)** Independent analysis of transcriptome (A) and protein (B) modalities from a CITE-seq dataset of cord blood mononuclear cells. Blue dot marks the same target cell in (A) and (B). Red dots denote the k=20 nearest neighbors to the target cell based on the transcriptome (A) or protein (B) modalities. **(C)** The RNA neighbors are averaged together to predict the molecular contents of the target cell, which can be compared to the actual measurements. Since the RNA neighbors represent a mixture of different T cell subsets, there is substantial error between predicted and measured protein expression levels for CD4 and CD8. **(D)** Same as in (C), but averaging protein neighbors. Since protein neighbors are all CD8 T cells, the predicted values are close to the actual measurements. We can therefore infer that for this target cell, the protein data is most useful for defining cell state, and assign it a higher protein modality weight. As described in Supplementary Methods, we perform the prediction and comparison steps in low-dimensional space. **(E)** We can integrate the modalities by constructing a Weighted Nearest Neighbor (WNN) graph, based on a weighted average of protein and RNA similarities. UMAP visualization and clustering of this graph. **(F)** Median RNA and protein modality weights for all cell types in the dataset. Modality weights were calculated for each cell without knowledge of cell type labels.

For each cell, we began by independently calculating sets of *k*=20 nearest neighbors for each modality. We found that for CD8^+^ T cells, the most similar RNA neighbors often reflected a mix of CD8^+^ and CD4^+^ T cells (in the RNA KNN graph, there are a total of 944 incorrect edges that connect CD8^+^ to CD4^+^ T cells). By contrast, protein neighbors were predominantly correctly identified as CD8^+^ T cells (in the protein KNN graph, 12 CD8^+^/CD4^+^ edges were identified). This reflects the particular utility of protein data when defining the state of these cells. Next, we independently averaged the molecular profiles of protein neighbors and RNA neighbors (i.e. predicted the molecular contents of a cell from its neighbors), and compared the averages to their original measured values. We found that for CD8^+^ T cells, protein KNN-based predictions were more accurate compared to RNA KNN-based predictions (Figure 1), while the converse was true for cDC (Figure 1C, D; Supplementary Figure 1).

We then leveraged the relative accuracy of these predictions to calculate RNA and protein modality “weights”, describing their relative information content for each individual cell. We provide a detailed mathematical description for each component of the WNN workflow in the Supplementary Methods, highlighting three key steps: 1) Obtaining within modality and cross-modality predictions, 2) Converting these to prediction affinities, based on a cell-specific bandwidth kernel, and 3) Calculating modality weights using a softmax transformation. The RNA and protein modality weights are non-negative, unique to each cell, and sum to 1.

Our final step integrates the modalities to create a ‘weighted nearest neighbor’ (WNN) graph. For each cell, we calculate a new set of *k*-nearest cells based on a metric which reflects the weighted average of normalized RNA and protein similarities (Supplementary Methods). The WNN graph is a single representation of a multimodal dataset, but should more accurately reflect the richness of both data types. For example, the WNN graph contained only 20 CD8^+^/CD4^+^ edges. Moreover, many common analytical tasks for single-cell data - including tSNE/UMAP visualization, clustering, and trajectory inference, can accept a user-specified neighbor graph as input. We therefore used our WNN graph to derive an integrated UMAP and clustering of our CITE-seq dataset (Figure 1E). In contrast to the separate analysis of either modality, our joint integration clearly separated CD4^+^ and CD8^+^ T cells, retained the identity of cDCs, and also uncovered additional sources of subtle heterogeneity within NK cells (Supplementary Figure 1). We observed that cells classified as CD8^+^ T cells were assigned higher protein modality weights, while DCs were assigned higher RNA modality weights, recapitulating our biological expectations despite the fact that the calculation of modality weights was unsupervised, and unaware of cell-type labels (Figure 1F).

### WNN analysis is a robust and flexible approach for multimodal analysis

We next further explored the performance of our WNN integration, assessed its robustness to fluctuations in data quality, and performed benchmarking against other recently developed methods. For these analyses, we used a more recently generated CITE-seq dataset of human bone marrow, representing 30,672 mononuclear cells with a panel of 25 antibodies. While the samples contained cells across the full spectrum of hematopoietic differentiation, the antibody panel was designed to separate groups of terminally differentiated cells.

Consistent with our previous example, WNN integration substantially increased our ability to resolve hematopoietic cell states (Figure 2A; Supplementary Figure 2). Once cell states were annotated through integrated multimodal clustering, we were able to discover differentially expressed (DE) genes and proteins in each group, further validating their biological identity and significance (Supplementary Figure 2). However, while these cell types are defined by both RNA and protein markers, the statistical power in unsupervised analysis of either modality separately was insufficient to identify these populations, demonstrating the importance of joint analysis. Indeed, when examining the cell-specific modality weights, we found that T-cell groups - and in particular, populations that were masked in scRNA-seq analyses - all received higher protein modality weights (Figure 2B).

**Figure 2:**
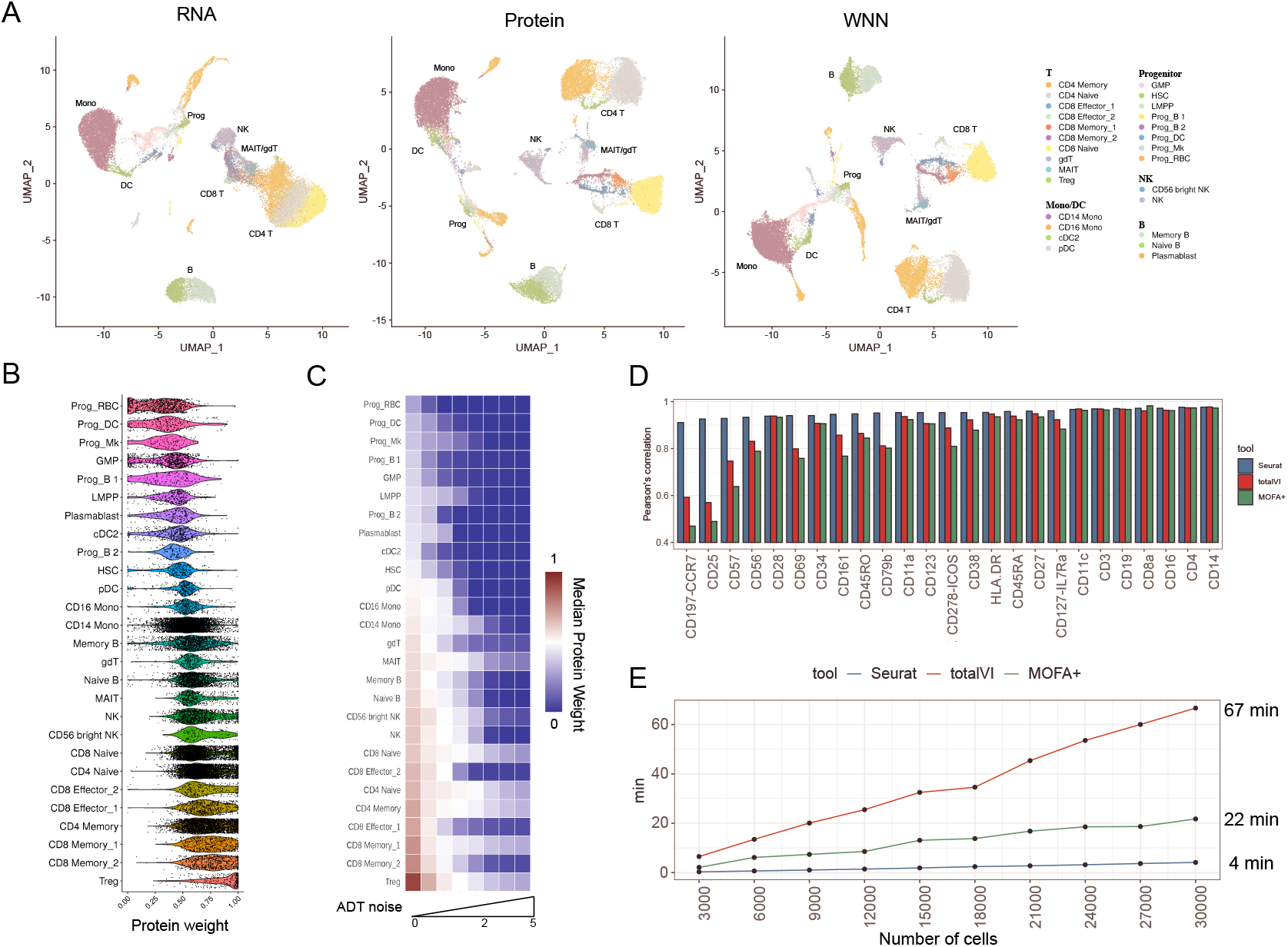
Benchmarking and robustness analysis for WNN integration. **(A)** Analysis of a CITE-seq dataset of human bone marrow mononuclear cells and 25 surface proteins. UMAP visualizations are computed using RNA, Protein, or WNN analysis. Cell annotations are derived from WNN analysis, and reveals heterogeneity within T cells and progenitors that cannot be discovered by either modality independently. Granular annotations, which more clearly indicate subpar performance when analyzing only one modality, are shown in Supplementary Figure 2. **(B)** Single-cell protein modality weights. Progenitor populations all receive low protein weights, while T cell populations receive high protein modality weights, consistent with the composition of the antibody panel which was tailored for differentiated cell types. **(C)** To test the robustness of WNN, we added increasing amounts of Gaussian noise to the protein data. Protein weights decrease to 0 in all cell types as noise levels increase. **(D, E)** Benchmarking WNN against totalVI and MOFA+. (D) The integrated latent space defined by WNN most accurately reconstructs expression levels for 25 proteins. (E) WNN analysis exhibits improved runtimes compared to competing methods. Additional benchmarking analyses in Supplementary Figure 2.

Conversely, each of the cell populations with the highest RNA weights represented hematopoietic progenitor populations. This was consistent with the composition of our protein panel, which did not contain surface markers distinguishing between groups of multipotent and lineage-committed CD34^+^ progenitors. As a result, our multimodal analysis was able to identify diverse populations of hematopoietic stem cells, lymphoid-primed multipotent progenitors (LMPP), and progenitors of erythroid, platelet, monocyte, B, and conventional/plasmacytoid DC lineages that could be recovered in scRNA-seq data, even though these groups lacked immunophenotypic markers in our CITE-seq experiment. We confirmed that varying the parameter *k* (set to 20 by default) across a range from 10 to 50 introduced only minor differences to the modality weights (Supplementary Figure 2), demonstrating that our results are robust to changes in this parameter.

These results suggest that integrated WNN analysis can provide necessary flexibility and allow one data type to compensate for weaknesses in another. We confirmed this using a simulation experiment, where we added increasing amounts of random Gaussian noise to the ADT data, in order to mimic increases in nonspecific binding (Figure 2C). We found that the increasing ADT noise led to a decrease in protein weights for all cell types, in a dose-dependent manner. Moreover, protein modality weights were assigned to 0 after a sufficient amount of protein noise was added, correctly instructing downstream analyses to focus only on scRNA-seq data.

We next benchmarked WNN analysis against two recently introduced methods for multimodal integration: Multi-omics factor analysis v2 (MOFA+) [25], which uses a statistical framework based on factor analysis, and totalVI [26], which combines deep neural networks with a hierarchical Bayesian model. Both methods integrate the modalities into a latent space, which we used to construct an integrated *k*-NN graph and a 2D UMAP visualization. We reasoned that we could quantify the performance of the different methods by comparing the similarity of each cell’s molecular state to its closest neighbors in the integrated latent space. We found that for each of the 25 proteins (Figure 2D), as well as the RNA transcriptome (Supplementary Figure 2), WNN analysis exhibited superior or equivalent performance to alternative approaches. The difference in performance was particularly striking for markers of regulatory (CD25) and effector (CD57) T cells. This was consistent with UMAP visualization, in which WNN was the only method where these populations were not intermixed with other groups (Supplementary Figure 2). WNN analysis also exhibited significant improvements in speed, ranging up to 15-fold when analyzing the full dataset (Figure 2E).

While we primarily demonstrate our approach on CITE-seq datasets, our strategy is applicable to diverse multimodal technologies. For example, recent developments have enabled the simultaneous measurement of ATAC-seq profiles and transcriptomes from single nuclei [12-14]. We applied WNN analysis to a dataset of 11,351 paired PBMC profiles generated by the 10x Genomics Multiome ATAC+RNA kit. We found that the combination of modalities exhibited maximal power to separate immune subsets (Supplementary Figure 3). Interestingly, similar to our CITE-seq analyses, we found that ATAC-seq data was more capable of separating naïve CD8^+^ and CD4^+^ T cell states due to reliable detection of cell type-specific open chromatin regions (Supplementary Figure 3). The separation of these clusters upon UMAP visualization (Supplementary Figure 3) was consistent with the number of incorrect naïve CD8^+^/CD4^+^ edges identified in each representation (RNA KNN: 984, ATAC KNN: 373, WNN: 322).

The combination of ATAC and RNA data also allowed us to identify differentially accessible DNA sequence motifs between our WNN-defined clusters. For example, we found that ATAC-seq peaks accessible in MAIT cells were highly enriched for motifs for the pro-inflammatory transcription factor RORγt [27, 28], which was also up-regulated transcriptionally in these cells (Supplementary Figure 3). We obtained highly concordant results when applying WNN analysis to ASAP-seq [29], a third multimodal technology, that pairs measurements of surface protein abundance with ATAC-seq profiles in single cells (Supplementary Figure 3). We conclude that WNN analysis is capable of sensitively and robustly characterizing populations that cannot be identified by a single modality, exhibits best-in-class performance, and can be flexibly applied to multiple data types for integrative and multimodal analysis.

### A multimodal atlas of the human peripheral blood mononuclear cells

While flow cytometry and CyTOF are widely used and powerful approaches for making high-dimensional measurements of protein expression in immune cells [30-33], CITE-seq’s use of distinct oligonucleotide barcode sequences provides a unique opportunity to profile very large panels of antibodies alongside cellular transcriptomes. In addition, we have recently demonstrated that the signal-to-noise for each antibody can be optimized for any individual marker as a function of antibody concentration, and shown that CITE-seq data quality does not deteriorate with greater amounts of total antibody [34]. We therefore curated and optimized a panel of TotalSeqA reagents encompassing 228 antibodies (Supplementary Table 1) comprising a diverse set of lineage and activation markers. One potential concern with large CITE-seq panels is that the signal from a small number of markers could saturate the overall library, drastically increasing the required sequencing depth. We developed a simple experimental procedure to address this (Supplementary Methods), first identifying 12 out of 228 markers that received a disproportionate number of reads in preliminary experiments, and then supplementing the panel with their unconjugated (barcode-free) antibodies to the panel in order to dampen the magnitude of their signal.

We leveraged the CITE-seq technology alongside our optimized antibody panel and integrative WNN analysis strategy to generate a multimodal atlas of human PBMC. We obtained PBMC samples from eight volunteers enrolled in an HIV vaccine trial [35, 36], with ages spanning from 20-49 years (median age of 36.5). For each subject, PBMCs were collected at three time points: immediately before (day 0), three days, and seven days following administration of a VSV-vectored HIV vaccine (Figure 3A). The total dataset consists of 24 samples and utilized ‘Cell Hashing’ [34] to minimize technical batch effects. For each sample, we profiled cells using 10X Chromium 3’ (using 228 TotalSeq A antibodies), representing a total of 161,764 cells (average of 8,003 unique RNA molecules/cell, 5,251 unique ADT/cell). We also profiled a total of 49,147 cells split across all samples using ECCITE-seq [24], which enables surface protein profiling with the 10X 5’ technology. While the latter experiments featured a smaller panel of 54 antibodies comprising a mix of lab-conjugated antibodies and TotalSeq-C reagents, reflecting availability of commercial conjugates at the time of the experiment, we were able to perform immune repertoire profiling of these cells as well. After NovaSeq sequencing, stringent quality control and doublet filtration (Supplementary Methods), our final dataset consists of 210,911 total cells, and allows us to profile cellular heterogeneity in both the resting (unvaccinated) and activated (post-vaccination) immune system.

**Figure 3:**
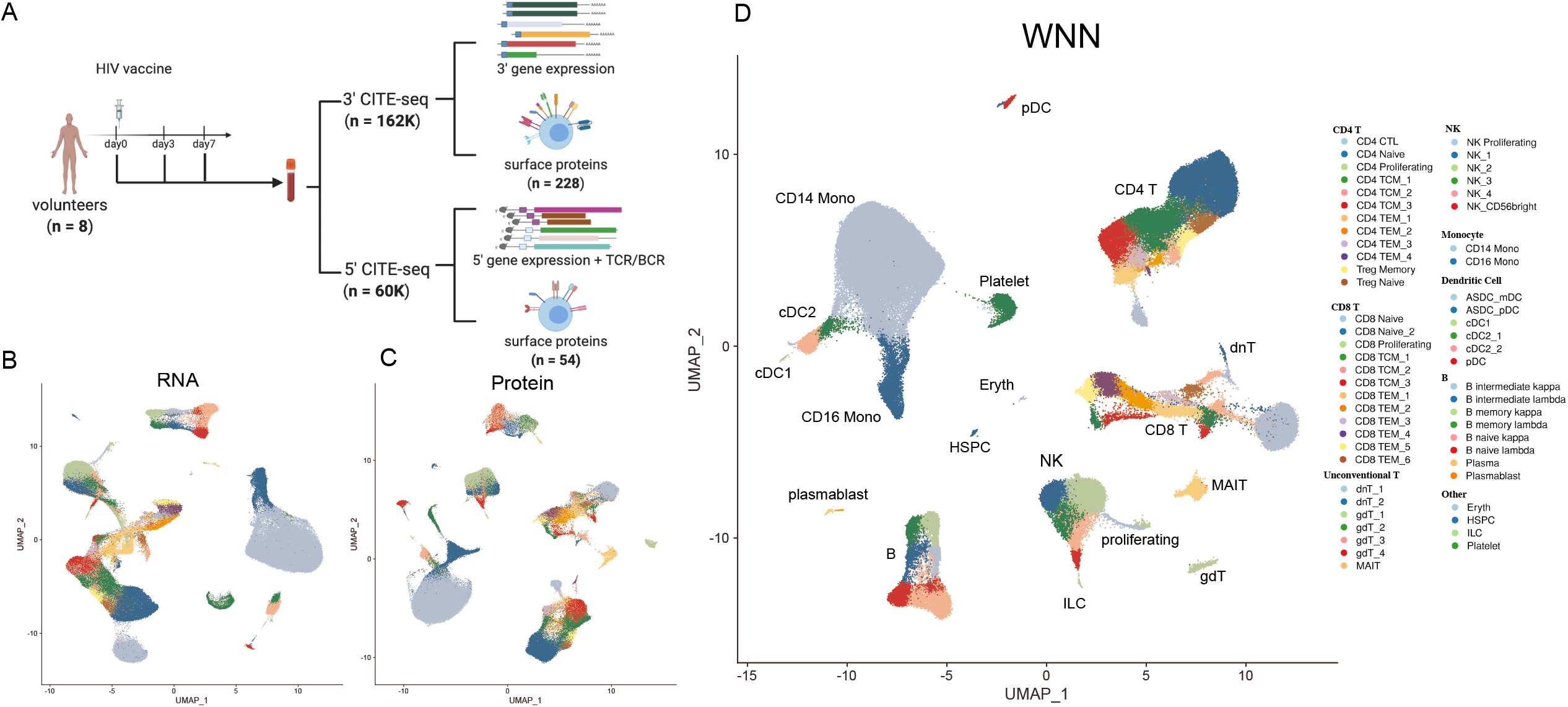
A multimodal atlas of human PBMC. **(A)** Experimental design schematic of the CITE-seq experiment. PBMC samples originate from eight volunteers pre (day 0) and post-vaccination (day 3 and day 7). We processed each sample with CITE-seq using the 10X 3’ (228 antibodies) and 10X 5’ (54 antibodies + BCR +TCR) technologies, yielding a total of 210,911 cells. **(B-D)** UMAP visualization of 161,764 cells 10X 3’ cells analyzed based on RNA data (B), protein data (C), or WNN analysis (D). Cell types were identified using unsupervised clustering of the WNN graph and grouped into three annotation tiers, ranging from eight broad categories, to 57 high-resolution clusters. UMAP visualization of 49,147 10X 5’ cells, mapped onto the 3’ reference data, is shown in Supplementary Figure 6.

We applied our ‘anchor-based’ workflow [37] to first integrate the samples together, enabling cells to cluster together based on their shared biological state, as opposed to sample-of-origin (Supplementary Methods). While this causes unvaccinated and vaccinated samples to cluster together initially, it enables us to annotate cell states consistently in all samples, and to learn cell-type specific responses in downstream analyses. We then performed joint analysis of both modalities using WNN integration, and as a comparative control, visualized the dataset using the RNA and protein modalities independently (Figure 3B-D).

We identified 57 clusters in WNN analysis, encapsulating all major and minor immune cell types, and revealing striking cellular diversity particularly within lymphoid lineages. With rare exceptions for infrequent cell types, each cluster included cells from all 24 samples. Our clusters could be readily grouped into larger categories, including CD4^+^ T cells (12 clusters), CD8^+^ T cells (12 clusters), unconventional T cells (7 clusters), NK cells (6 clusters), B cells, plasma cells, and plasmablasts (8 clusters), dendritic cells and monocytes (8 clusters), and rare clusters of hematopoietic progenitors, platelets, erythrocytes and circulating innate lymphoid cells (ILC). To assist in the interpretation of our clusters, we assign each cell three annotations with increasing granularity (Level 1, 8 categories; Level 2, 30 categories; Level 3; 57 categories). While we saw the greatest level of heterogeneity within T cell subsets, our analysis clearly identified heterogeneous subsets of myeloid cells that were fully concordant with recent high-resolution scRNA-seq analyses of sorted populations, including extremely rare populations (0.02%) of dendritic cells defined by the expression of *AXL* and *SIGLEC6* [38, 39] (ASDC; Supplementary Figure 4). We also identified substantial heterogeneity in the expression of inflammatory genes such as *IL1B* and *CCL3* within monocyte populations, but as this heterogeneity varied across different volunteers, we conservatively did not further subdivide these states (Supplementary Figure 4).

We next identified differentially expressed RNA and immunophenotype markers for each cluster. We found that each cluster exhibited distinct molecular patterns and biomarkers for both modalities (heatmap for CD4^+^ and CD8^+^ T cell states in Figure 4A; additional heatmaps in Supplementary Figure 5). Moreover, these identified biomarkers were invariant across human volunteers and vaccination timepoints. Observing consistent cluster biomarkers in multiple samples provides evidence that our observed heterogeneity is biologically reproducible, and not driven by overclustering. Despite the fact that clusters were enriched for both RNA and protein markers, our ability to identify these groups was substantially reduced without WNN analysis, as multiple clusters blended together when performing separate analysis of either RNA or protein data (Figure 3B, C). We conclude that multimodal integration is essential for the unsupervised discovery and annotation of immune cell states, but that once these states are enumerated, supervised differential analyses are capable of sensitively describing markers that define their molecular state.

**Figure 4:**
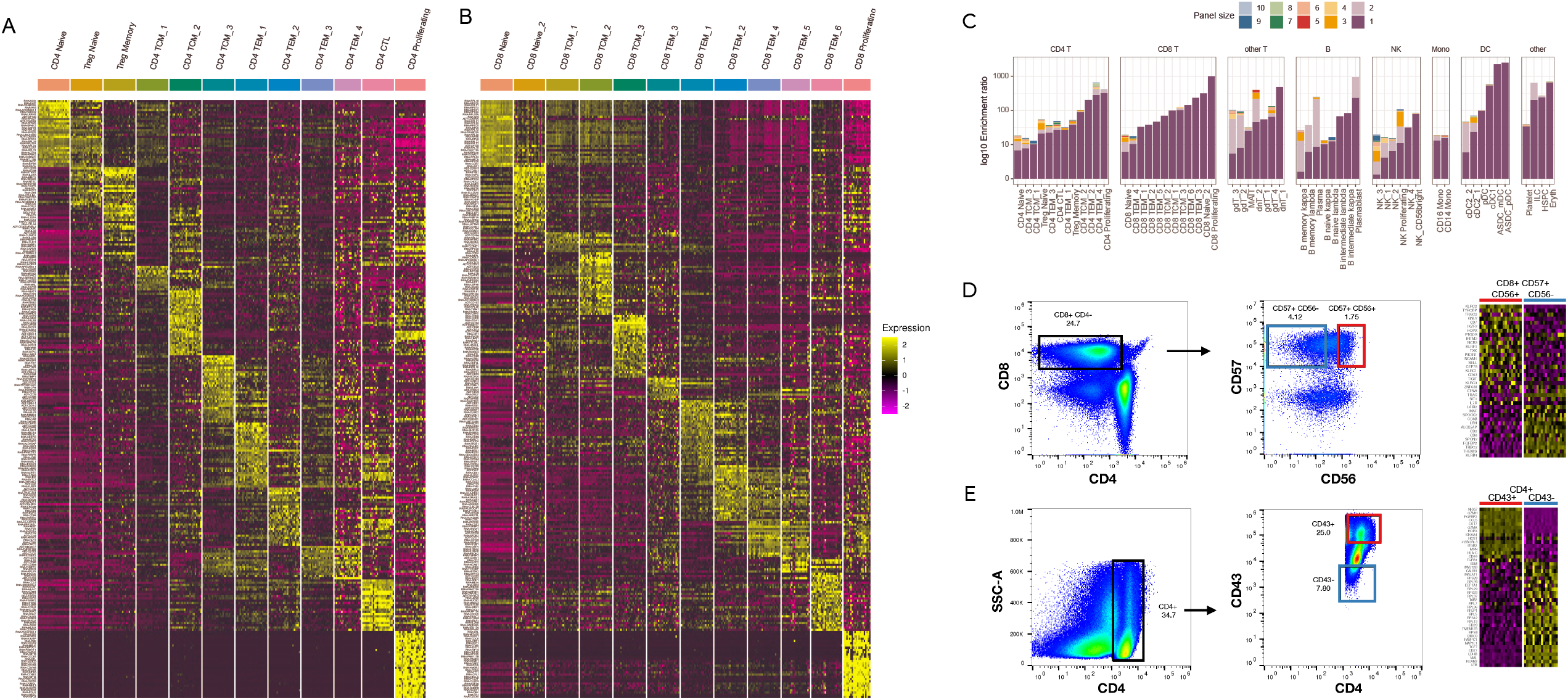
Multimodal biomarkers of immune cell states. **(A)** Heatmap of CD4^+^ T cell states. Markers include the best RNA and protein features identified by differential expression (DE). Heatmap displays pseudobulk averages where cells are grouped by cell type, donor, and vaccination time point and demonstrates that markers do not vary across different PBMC samples. **(B)** Same as in (A) but for CD8^+^ T cell states. Additional heatmaps are shown in Supplementary Figure 5. **(C)** For each of our 57 clusters, we calculated the optimal surface marker enrichment panels based on our CITE-seq data. Barplots show the ability of the panels to enrich for each cell type *in silico.* The composition of each panel is shown in Supplementary Table 2. **(D)** Validation of predicted marker panels for the CD8_TEM_5 cluster. We sorted cells based on the marker panels identified in (C), and performed bulk RNA-seq. Heatmap is ordered by genes expected to be DE based on our CITE-seq dataset, and are validated by bulk RNA-seq. **(E)** Same as in (D) but for CD4 CTL cells.

Due to the robust detection of protein features in CITE-seq combined with the size of our antibody panel, we reasoned that we could discover small panels of immunophenotypic markers for each cluster that could be used, for example, to perform targeted enrichment through flow cytometry. We used stepwise variable selection coupled with logistic regression (Supplementary Methods) to identify the best antibody marker panels of different sizes (1-10 markers) for each subset, and calculated the level of enrichment *in silico* (Figure 4C). We found that a single marker was capable of achieving effective enrichment of at least ten-fold for 45 clusters, while a panel with three markers was sufficient to achieve 10-fold enrichment for 55 clusters.

We confirmed that this marker discovery procedure identifies effective panels for well-characterized populations (plasmacytoid DC (pDC): CD123^+^, MAIT cells: CD3^+^ CD161^+^ TCRvα7.2^+^, CD4 Naive: CD4^+^ CD45RA^+^ CD45RB^+^). In other instances, for example, cytotoxic populations of CD4^+^ lymphocytes, our analysis identified CD43 as a marker with high enrichment power that has not been previously reported. For this population, as well as a subgroup of highly cytotoxic CD8^+^ T cells (CD8_TEM_5), we successfully validated our enrichment panels in an independent set of PBMCs from healthy donors by conventional flow cytometry followed by bulk RNA-seq (Supplementary Methods). In both cases we examined the expression level of genes that we expected to be DE based on our CITE-seq data, and we observed clear agreement between the sorted profiles and CITE-seq clusters (Figure 4D, E). Notably, our flow cytometry experiments utilized the exact antibody clones represented in the CITE-seq experiment, which can help to ensure that the two assays will return concordant results. We report each of these panels in Supplementary Table 2 to facilitate similar experiments for additional clusters in our dataset. We note that while these panels can achieve high levels of enrichment, even optimally sorted groups may contain a minority of contaminating cells from other states. We show precision and recall metrics for each panel in Supplementary Figure 4, demonstrating that it remains challenging to sort truly ‘homogeneous’ populations of high-resolution subsets using a small number of markers.

### Multimodal heterogeneity within lymphoid populations

Our integrated WNN analysis reveals a rich diversity of T cell states that is not typically captured in scRNA-seq analyses, including CD4^+^ regulatory T cells, MAIT cells, multiple subpopulations of γδ and double-negative T cells, along with heterogeneous subpopulations of naive, memory, and effector states. Even for populations that have been previously described by flow cytometry or CyTOF, our dataset represents a powerful resource to characterize enriched RNA and protein biomarkers for each cell state. However, we also observed sources of heterogeneity that have not been previously well-characterized in peripheral blood.

For example, within CD8^+^ memory T cells, we identified distinct subpopulations defined by bimodal and mutually exclusive expression of the integrin proteins CD49a and CD103 (Figure 5A). While we identified these cells in peripheral blood, expression of these proteins has traditionally been strongly associated with tissue-resident memory (TRM) cells, where integrins help mediate adhesion to epithelial cells or the extracellular matrix [40, 41]. CD8^+^ CD103^+^ T cells expressed high surface protein levels of the heterodimeric co-binding partner integrin beta-7 (Figure 5B), while expression was absent in CD8^+^ CD49a^+^ groups. We validated the presence of the populations in independent healthy PBMC samples by performing flow cytometry for the same markers (Figure 5C, D). In addition, we identified modules of differentially expressed genes between these two groups (Figure 5E), which were enriched for T cell activation, differentiation, signaling response, and chemotaxis modules (Figure 5F). Both populations did not express the canonical resident marker CD69 [42, 43] (Supplementary Figure 6), suggesting that they are not TRMs that have temporarily detached and re-entered circulation. Instead, these subpopulations may represent cells that are preparing to become tissue-resident and have already begun to acquire distinguishing molecular characteristics.

**Figure 5:**
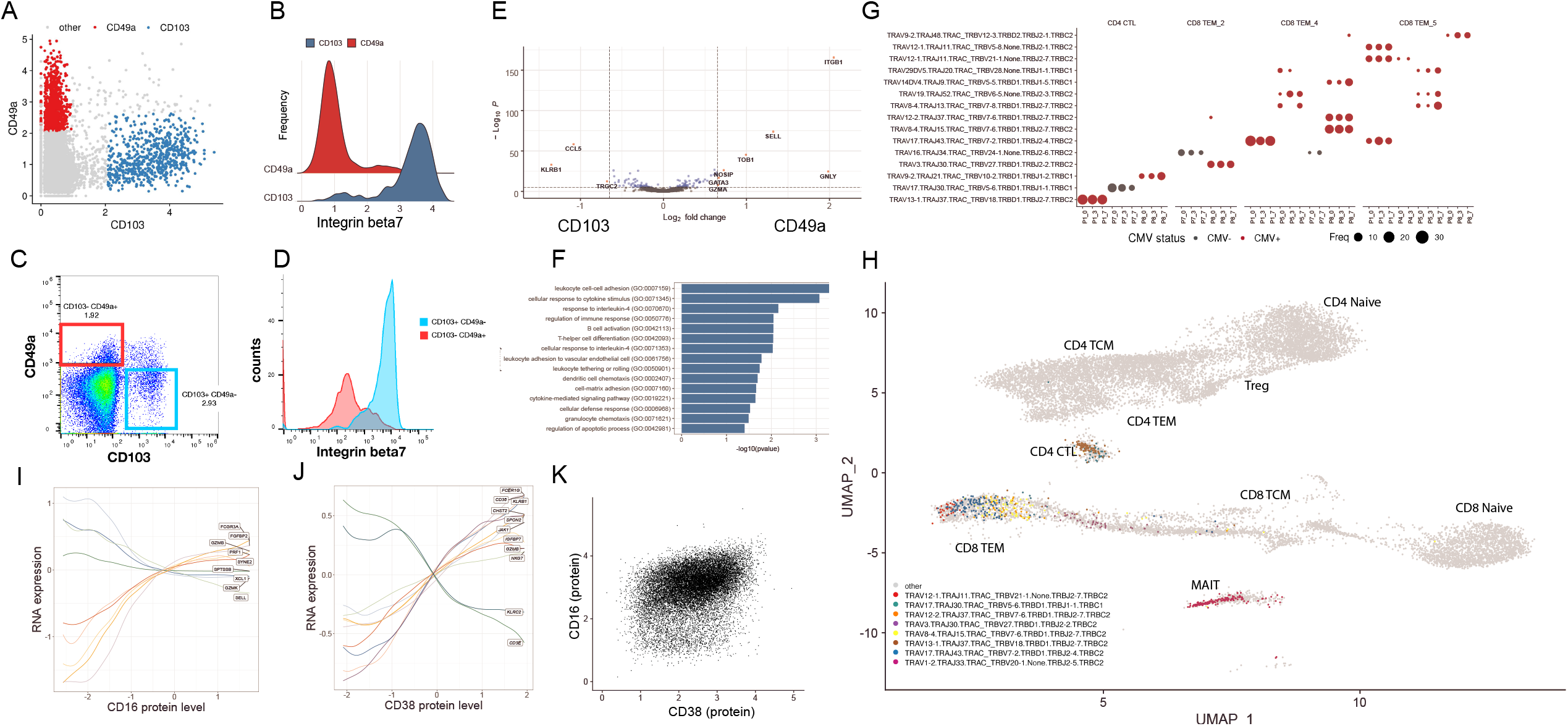
Characterizing heterogeneity within lymphoid populations. **(A)** Mutually exclusive expression of the integrin proteins CD103 and CD49a within CD8+ T memory cells, as measured by CITE-seq. **(B)** Differential expression of Integrin-7 between CD103^+^ CD49a^-^ and CD103^+^ CD49+ populations as measured by CITE-seq. **(C-D)** Flow cytometry validates the presence of these populations. Plots are the same as in (A-B), but generated via flow cytometry. **(E-F)** Differentially expressed genes, and enriched gene ontology terms, between CD103^+^ CD49a^-^ and CD103^-^ CD49+ populations. **(G)** Dot plot showing the representation of the fifteen most abundant T cell clonotypes in the dataset. For space, only the VDJ regions are shown on the y-axis, but all cells in a clone share identical CDR3 sequences. Clones reside in a restricted set of cytotoxic and effector cell states, and are shared across vaccination time points. Size of each dot represents the number of cells in the clonotype. Clones present in donors who were classified as CMV-positive are colored in red. **(H)** Cells within a clone exhibit similar molecular profiles. Grey dots represent T cells where TCR sequence was measured using the 10X 5’ assay. Cells from the eight most highly represented clonotypes are highlighted as colored dots.**(I-K)** Heterogeneity in NK cells is defined by two gradients correlating with CD16 and CD38 protein expression. **(I)** NK cells are ordered by their quantitative expression of CD16 protein expression. Rolling averages for the expression of genes that correlate positively or negatively with CD16 are shown as smoothed lines. **(J)** same as (I) but for CD38. **(K)** CD38 and CD16 protein expression define two separate gradients, and are uncorrelated in NK cells.

In addition to characterizing heterogeneity in mRNA and protein expression, we leveraged our 5’ dataset to explore the relationship between molecular state and TCR sequence. We obtained productive TCR α/β sequences representing 16,060 distinct clones, where all cells within a clone share the exact same CDR3α and CDR3β sequences. Overall clonal diversity was consistent across vaccination timepoints, consistent with an expected lack of a lymphoid response to vaccination within 7 days, and 97% of clones consisted only of a single cell. However, we also observed the presence of expanded clonal populations. As a positive control, we observed populations with highly restricted usage of TCRa sequences: both MAIT and invariant NKT cells exhibited closely related transcriptional profiles [44] and semi-invariant repertoires across multiple volunteers (Supplementary Figure 6).

Excluding these populations, we identified 31 additional expanded clones consisting of at least 10 cells (Figure 5G). In each case, cells within a clonal population exhibited extremely similar molecular profiles (Figure 5H), representing subgroups of CD8^+^ T cells (primarily within our previously identified CD8_TEM_4 and CD 8_TEM_5 clusters), as well as cytotoxic CD4^+^ T cells (CD4 CTL). Each clone typically represented cells from a single volunteer, but could be independently found across multiple timepoints, including before vaccination (Figure 5G). As our sample volunteers were generally middle-aged and otherwise healthy, we considered the possibility that overexpanded clones could be related to Cytomegalovirus (CMV) infection [45]. We assessed the CMV status of each volunteer by stimulating PBMC with a CMV peptide pool and performing intracellular cytokine staining to determine responses in CD8^+^ T cells (Supplementary Methods, Supplementary Table 3), identifying five positive and three negative volunteers. We found that the five positive samples accounted for 91% of cells within expanded clones.

We note that while WNN integration improves the ability to discover distinct cell subpopulations, it can also improve the characterization of cellular trajectories and continuous sources of heterogeneity. For example, within B cells, we identified a continuous trajectory connecting naïve to memory cells defined by the canonical protein markers IgD and CD27, along with a module of correlated genes (Supplementary Figure 6). Similarly, NK cells were subdivided into five clusters, representing variation across a continuous landscape. Our data shows that the traditional division of NK cells into CD56-bright and CD56-dim categories represents a broader continuum defined by CD16 expression, alongside a module of genes that modulate cytotoxicity and correlate both positively and negatively with this marker (Figure 5I).

We also observed a second gradient defined by CD38 expression, that to our knowledge has not been previously described. Notably, *KLRC2*, which encodes the NK activating receptor NKG2C was negatively associated with this continuum, while the signaling adapter *FCER1G* was positively associated (Figure 5J). This expression pattern is consistent with the development of ‘adaptive’ or ‘memory-like’ NK cells observed in CMV seropositive individuals [46, 47]. Notably, we observed consistent trends when restricting our analysis only to individuals with either positive or negative CMV T cell responses (Supplementary Figure 6). We also observed consistent results in an independent CITE-seq dataset of human PBMC [48]. Our results indicate that this phenotype does not represent a strictly binary phenomenon and may not be specific to CMV response. Finally, we observed minimal correlation between CD38 and CD16 expression (Figure 5K), demonstrating that NK cells fall along a two-dimensional gradient defined by these markers.

Taken together, these results demonstrate that our dataset represents a powerful resource to enumerate cell states in the immune system, identify optimal reagents for cell-type specific enrichment, and to understand the molecular heterogeneity in clonally related or antigen-specific cell groups. As these results are consistent in both pre and post-vaccination timepoints, they likely describe general characteristics of the healthy immune system. We have released a web portal at https://atlas.fredhutch.org/nygc/multimodal-pbmc to facilitate community exploration of this resource.

### Characterizing the initial innate response to vaccination

We next explored our dataset to characterize the response to vaccination for each of our previously identified cell types. We were particularly interested to identify cell populations that contribute most strongly to the innate immune response, which is expected to be highly activated at our first vaccinated time point (day 3), and subsequently dampen in our second time point (day 7) as seen with another non-replicating viral vectored HIV vaccine [49]. We were interested in responses that manifested as cell-type specific changes either in gene expression, protein expression, or abundance across vaccination timepoints.

In order to quantify strength of response for each cluster, we first identified the number of differentially expressed genes and proteins between pre-vaccination and day 3 cells for each cell type. However, as pergene tests are highly sensitive to the number of cells in each cluster, we also leveraged an alternative approach (‘perturbation score’, as described in [50]), which retains the statistical power to distinguish cells from different timepoints on the basis of the full transcriptome. As expected, we observed robust responses in a subset of myeloid subpopulations, but only minimal responses in lymphoid groups (Figure 6A, B). Response patterns were also largely consistent across samples, with the exception of one volunteer exhibiting a highly activated immune system in advance of vaccination, that was removed from further analysis (Supplementary Figure 7).

**Figure 6:**
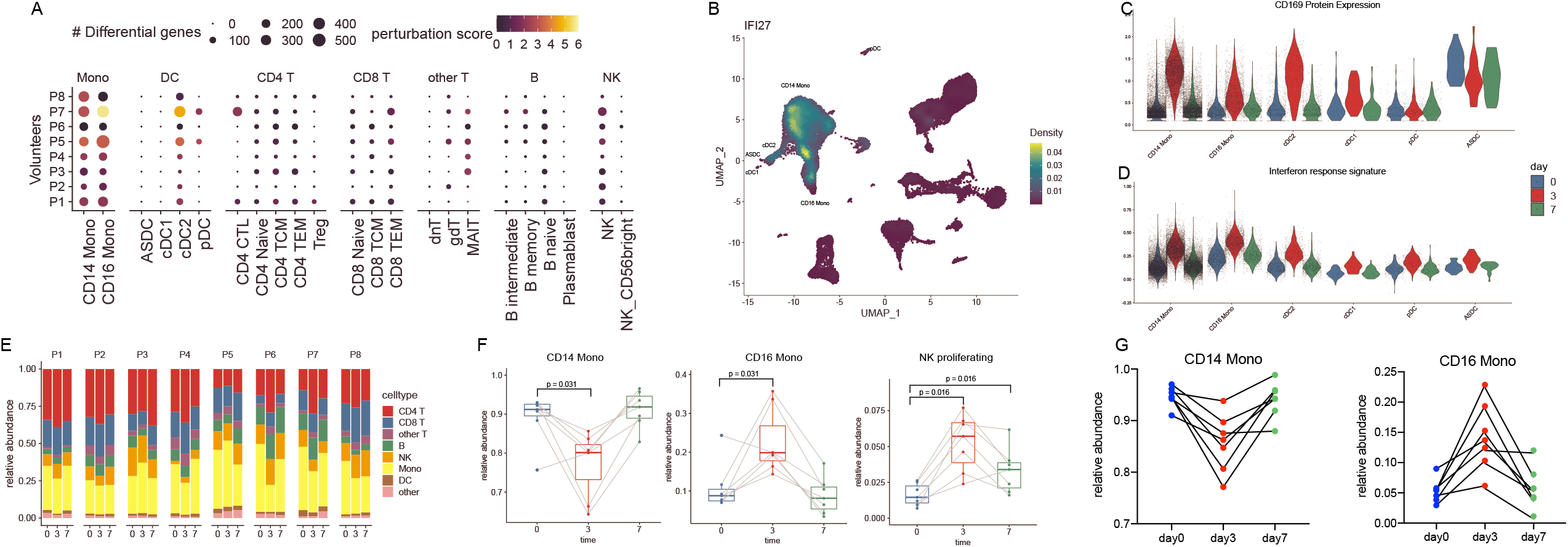
Identifying cell-type specific responses to vaccination. **(A)** For each of our Level 2 annotated cell clusters, we calculated the number of differentially expressed genes between unvaccinated (day 0) and day 3 samples (size of each dot). As each per-gene test is highly sensitive to the number of cells, we also calculated a ‘perturbation score’, which reflects the strength of the molecular response based on the whole transcriptome (color of each dot). **(B)** Density plot, produced by the Nebulosa package, showing the expression of canonical interferon response gene IFI27 **(C)** Violin plot showing the protein up-regulation of Siglec-1 (CD169) in single cells from day 3 samples, along with a signature of interferon response **(D)**, in select cell types. In (A-D) we consistently observe robust responses only in CD14+ monocytes, CD16+ monocytes, and cDC2 DC. **(E)** Barplot showing that the frequency of broad groups (Level 1 annotations) is stable across the vaccination time course. **(F)** Within these broad categories, the relative abundance of classical monocytes, nonclassical monocytes, and proliferating NK cells across the vaccination time course. p-values are computed using a paired Wilcoxon test. **(G)** Relative abundance of monocyte populations as measured by flow cytometry.

We observed the strongest changes in both CD14^+^ classical and CD16^+^ non-classical monocytes, as both cell types up-regulated a shared module of 62 genes highly enriched for transcripts responsive to type I interferon (Figure 6A, B; Supplementary Figure 7. Visualization in 6B from [51]). In addition, we identified Siglec-1 (CD169) as a protein response biomarker that was robustly induced only in day 3 samples (Figure 6C). When we examined dendritic cell populations, we observed a similarly robust response only within CD1C^+^ cDC2 cells. Contrastingly, closely related populations of CD141^+^ cDC1, as well as ASDC and pDC, exhibited minimal responses, and we did not detect any DE genes before and after vaccination for these groups (Figure 6A). This indicates that within DC subgroups, cDC2s may perform an important role in the downstream priming and activation of the adaptive immune system during this vaccine response.

We next attempted to identify cell types exhibiting changes in abundance during our time course. We did not observe significant changes in the overall abundance of broad immune classes (Figure 6E; level 1 annotations), so we focused on identifying more subtle compositional changes within these broader groups. For example, while the overall proportion of monocytes was consistent across timepoints, there was a strong shift in the ratio between classical and non-classical populations between day 0 and day 3 (Figure 6F). We validated this result, as well as the observed return to baseline ratios at day 7, using flow cytometry on the same samples (Figure 6G). We did not observe changes within lymphoid cells with one exception: a small population of NK cells expressing proliferation and cell cycle genes (NK_proliferation), consistently increased upon vaccination (Figure 6F). These findings were reproducible in independent analyses of the 3’ and 5’ scRNA-seq experiments and persisted in both day 3 and day 7 samples (Figure 6F; Supplementary Figure 7). This finding may reflect an early step in the development and maturation of NK cells, a key component of the NK cell-mediated innate immune response [52].

### Mapping query datasets to multimodal references

Single-cell transcriptomic profiling of the immune system has become routine, not only for healthy subjects, but also in multiple clinical contexts including for patients hospitalized with COVID-19. These datasets are typically processed using a workflow that consists of unsupervised clustering and ensuing annotation of cell types. This process assumes minimal prior knowledge and is ideally suited for cell type discovery. However, having constructed a multimodal reference of the immune system, we sought to leverage this dataset to assist in the analysis and interpretation of additional single-cell experiments profiling human PBMC (queries), even if only the transcriptome was profiled.

We first apply a procedure known as ‘supervised principal component analysis (sPCA) [53] to the transcriptome measurements in our reference dataset. Instead of seeking to identify a low-dimensional projection that maximizes total variance as in PCA, sPCA identifies a projection of the transcriptome dataset that maximally captures the structure defined in the WNN graph. Formally, given a gene expression matrix X and a WNN graph Y, sPCA identifies the transformation matrix U that maximizes the Hilbert-Schmidt Independence Criterion measure between a linear kernel of U^T^X and Y (Supplementary Methods). Informally, sPCA allows the weighted transcriptome and protein measurements to help ‘supervise’ the analysis of gene expression data and identify the optimal transcriptomic vectors (gene modules) that define the cell states in our multimodal reference.

We compute this sPCA transformation on our reference (where both mRNA and protein were measured simultaneously), but can subsequently rapidly project this transformation onto any scRNA-seq query dataset. Combining this transformation with our previously described ‘anchor’-based framework [37] allows us to place each scRNA-seq query cell on the previously defined reference UMAP visualization (Supplementary Methods), and to annotate its identity based on reference clusters. We note that in contrast to our previously developed scRNA-seq integration algorithms [37, 54], the reference dataset and visualization can remain constant during this procedure.

We found that this supervised mapping procedure dramatically improved our ability to analyze and interpret query scRNA-seq datasets compared to unsupervised analysis. We examined a recently generated dataset of human PBMC prior to flu vaccination, which measured the transcriptomes of 53,099 cells alongside 82 surface proteins. We mapped this dataset onto our reference using only the transcriptome data and transferred our Level 2 annotations, revealing the presence of multiple high-resolution lymphoid subsets (Supplementary Figure 8). We verified the accuracy of our predictions using the query protein data, which was held out of the reference mapping procedure, yet revealed expression patterns based on our predicted annotations that were fully concordant with our reference dataset. For example, cells that were annotated as regulatory T cells expressed CD25^+^ in the CITE-seq data, and we observed similar results for MAIT cells (CD161^+^), memory (CD45RA^-^ CD45RO^+^) and naive (CD45RA^+^ CD45RO^-^) T cells, and circulating ILC (CD117^+^ CD25^+^, Supplementary Figure 8). We benchmarked our method against scArches, a recently developed method for mapping scRNA-seq queries to reference datasets [55] and observed that our approach yielded substantial improvements in accuracy and performance (Figure 7A, B; Supplementary Figure 8).

**Figure 7:**
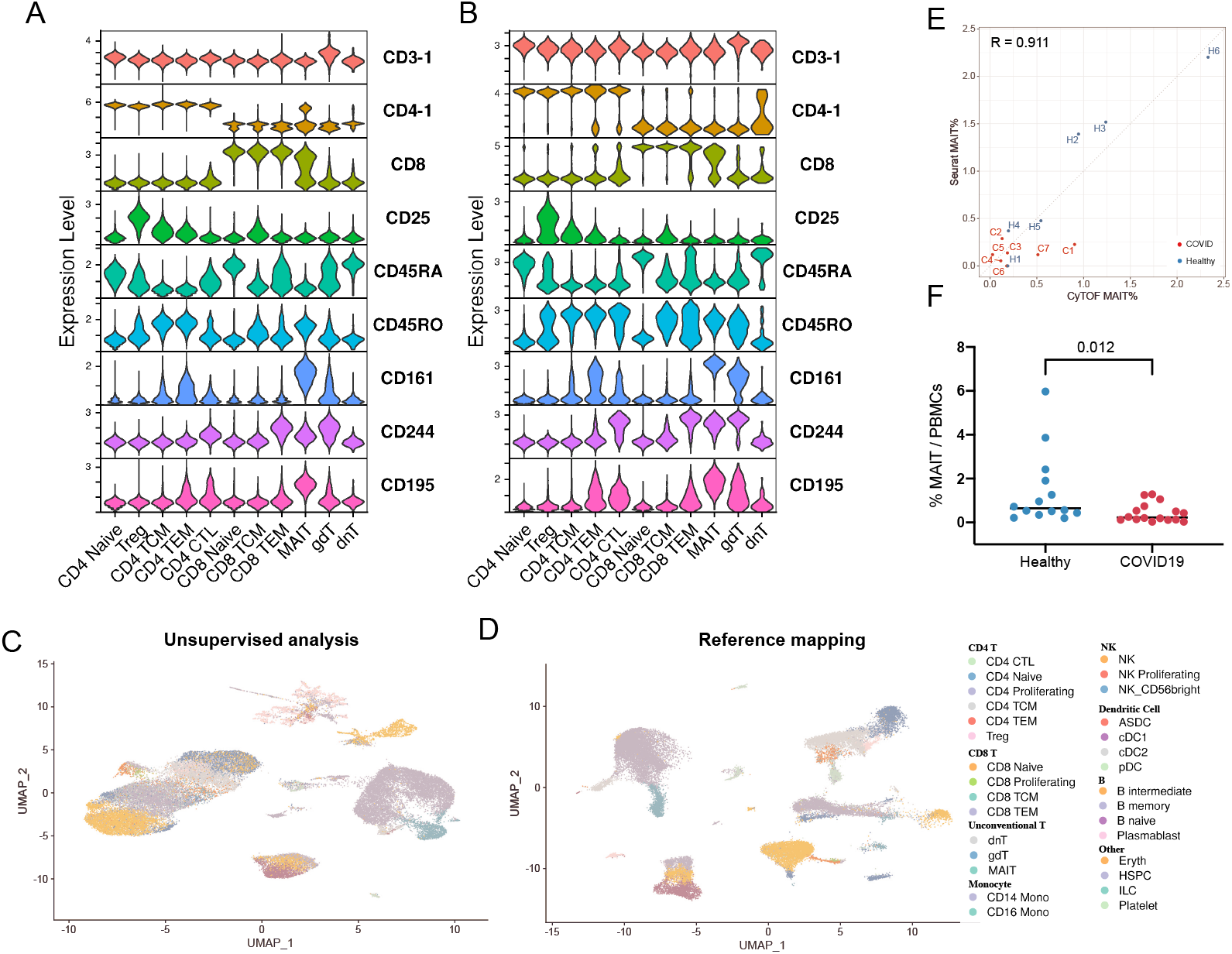
Supervised mapping of immune perturbations. **(A)** Violin plots showing the expression patterns for nine proteins in our CITE-seq dataset. Cells are grouped by their WNN-defined T cell Level 2 annotations **(B)** Violin plots for the same proteins in an independent CITE-seq dataset of human PBMC (Kotliarov et al., 2020). Cells are grouped based on their predicted annotations from transcriptome-based reference mapping. The protein data was withheld from the mapping, but displays the same patterns as in (A). **(C)** UMAP visualization of (Wilk et al, 2020) scRNA-seq dataset, which includes 44,721 PBMC from patients hospitalized with COVID-19, and healthy controls. UMAP was computed using unsupervised analysis. **(D)** Same as in (C), but after the dataset has been mapped onto our multimodal reference. Cells are colored by their predicted level-2 annotations. **(E)** Quantification of MAIT cell abundance based on scRNA-seq reference mapping (y-axis), and CyTOF (x-axis), for the samples in (Wilk et al, 2020). The Pearson correlation between these two methods are 0.911. **(F)** CyTOF quantification of MAIT cell abundance in PBMC samples from COVID-19 patients, and healthy controls. p-values are computed using a unpaired Wilcoxon test.

We next applied our mapping approach to a recent scRNA-seq study analyzing PBMC samples from seven patients hospitalized with COVID-19, alongside six healthy controls [56]. The original publication performed unsupervised clustering on the full dataset and identified six T cell clusters (three CD4^+^ T, two CD8^+^ T, and γδ T cells). In our supervised analysis we transferred our Level 2 annotations, successfully dividing T cells into the 12 groups (Figure 7C, D). Notably, populations of developing and differentiated neutrophils, which were identified by the original manuscript as being uniquely present in COVID-19 samples but were absent from our reference, could not be successfully mapped (Supplementary Figure 8).

We leveraged our supervised annotations to test for differences in cell type abundance across disease conditions. Our findings recapitulated the original unsupervised analysis, for example, highlighting increases in plasmablast frequency during COVID-19 response (Supplementary Figure 8). However, we also observed proportional shifts in cell states that were not detected in unsupervised clustering, but were successfully annotated after reference mapping. In particular, we observed a depletion of MAIT cells in COVID-19 samples compared to healthy controls. To validate our findings, we performed CyTOF on both the original samples and a validation cohort of 16 additional samples. We observed strong quantitative agreement (R=0.911) in the fraction of MAIT cells predicted by scRNA-seq and measured by CyTOF in the original cohort (Figure 7E). Moreover, CyTOF analysis of the larger sample set identified a depletion of MAIT cells in COVID-19 samples (Figure 7F; Supplementary Figure 8). This change in abundance may reflect these cells exiting circulation to play protective roles in barrier tissues during the antiviral immune response [57–59].

## Discussion

In order to leverage multiple data types to define cellular identity, we developed Weighted Nearest Neighbor analysis, a computational method that learns the information content of each modality and generates an integrated representation of multimodal data. By calculating cell-specific modality weights, WNN analysis solves an important technical challenge for the analysis of multimodal datasets and allows for flexible application across a range of modalities and data types. We demonstrate throughout this manuscript that performing downstream analyses on a weighted combination of data types dramatically improves our ability to characterize cellular diversity.

We apply our approach to analyze a dataset of human PBMC featuring paired transcriptomes and measurements of 228 surface proteins, representing a multimodal atlas of the immune system. We leverage this resource to characterize extensive lymphoid heterogeneity that has not been previously observed by scRNA-seq alone, including the heterogeneous expression of integrin proteins on circulating memory T cells, a gradient of adaptive-like responses in NK cells, and tightly clustered clonal populations within effector and cytotoxic groups. Our data also enables us to explore the response of the innate immune system to vaccination, highlighting specific response biomarkers, as well as the heterogeneous responses of conventional DCs. Importantly, we demonstrate that CITE-seq data can be easily mined to identify the best immunophenotypic marker panels for any subpopulation of interest. These marker panels can be used for flow cytometry with the same antibody clones in our CITE-seq panel, facilitating rapid enrichment and downstream analysis of these groups, and broadening the value of our resource.

In addition to constructing a multimodal reference, we demonstrate the ability to map scRNA-seq data onto this dataset. We accomplish this via a supervised version of principal component analysis to identify the best transcriptomic modules which delineate our WNN-defined cell types. Supervised mapping represents an attractive alternative to unsupervised analysis, and we show how this workflow can improve cell type identification and robustly integrate samples from multiple donors and disease states. To assist the community in utilizing our resource, we have created a web application, freely available at http://www.satijalab.org/azimuth, which enables users to rapidly map their own datasets online, automating the process of visualization and annotation. Using this approach, a dataset of 50,000 cells can be fully processed and mapped in less than five minutes. As the profiling of human PBMC under a variety of disease states becomes increasingly routine, the ability to perform automated mapping of these datasets will facilitate the characterization of complex immune responses, and the discovery of pathogenic populations.

Lastly, we note that the modality weights learned in our procedure serve not only as a proxy for the technical quality of a measurement type, but may also reflect the biological importance of each modality in determining cellular identity. For example, our analyses of human bone marrow demonstrated that progenitor cells and differentiated cells exhibited divergent modality weights. As future technologies enable the simultaneous measurement of modalities spanning the central dogma including chromatin state, DNA methylation, transcription, lineage, spatial location, and protein levels - WNN analysis may help to reveal how subpopulations of cells differentially utilize these modalities to regulate their current state and future potential. While our current implementation of WNN analysis focuses on the analysis of two modalities, the framework can be easily extended to handle an arbitrary number of simultaneous measurements as these technologies mature. Integrative multimodal analysis therefore provides a path forward to move beyond the partial and transcriptome-focused view of a cell, and towards a unified definition of cellular behavior, identity, and function.

## Data Availability

Seurat v4 is released under the open source GPLv3 license, and all code is available at www.github.com/satijalab/seurat. Installation instructions, tutorials, and documentation are available at www.satijalab.org/seurat

CITE-seq data generated for this manuscript is available to download and explore at https://atlas.fredhutch.org/nygc/multimodal-pbmc/

To facilitate the mapping of new query datasets to the multimodal PBMC reference described in this manuscript, we have released an automated web app, Azimuth: www.satijalab.org/azimuth

## Acknowledgements

We acknowledge Dan Littman and members of the Satija and Technology Innovation Labs for general discussion. This work was supported by the Chan Zuckerberg Initiative (EOSS-0000000082 to RS, HCA-A-1704-01895 to PS and RS), the National Institutes of Health (RM1HG011014-01 to PS and RS, 1OT2OD026673-01to RS, DP2HG009623-01 to RS, R21HG009748-03 to PS, U19AI128914 to RG, UM1 AI068618 to JMM), the Bill and Melinda Gates Foundation (OPP1113682 to CAB and AJR), and the Brotman Baty Institute (MZ).

## Author contributions

PS and RS conceived the study. YH, SZ, AB, CD, MZ, PH, JJ, AS, TS performed computational work, supervised by RG and RS. SH, EAN, WMM, MJL, AJW, MS, EP, EPM, LBF, BY, AJR performed experimental work, supervised by JME, CAB, and PS. All authors participated in interpretation and writing the manuscript.

