## Supplementary Figures for "Integrated analysis of multimodal single-cell data"

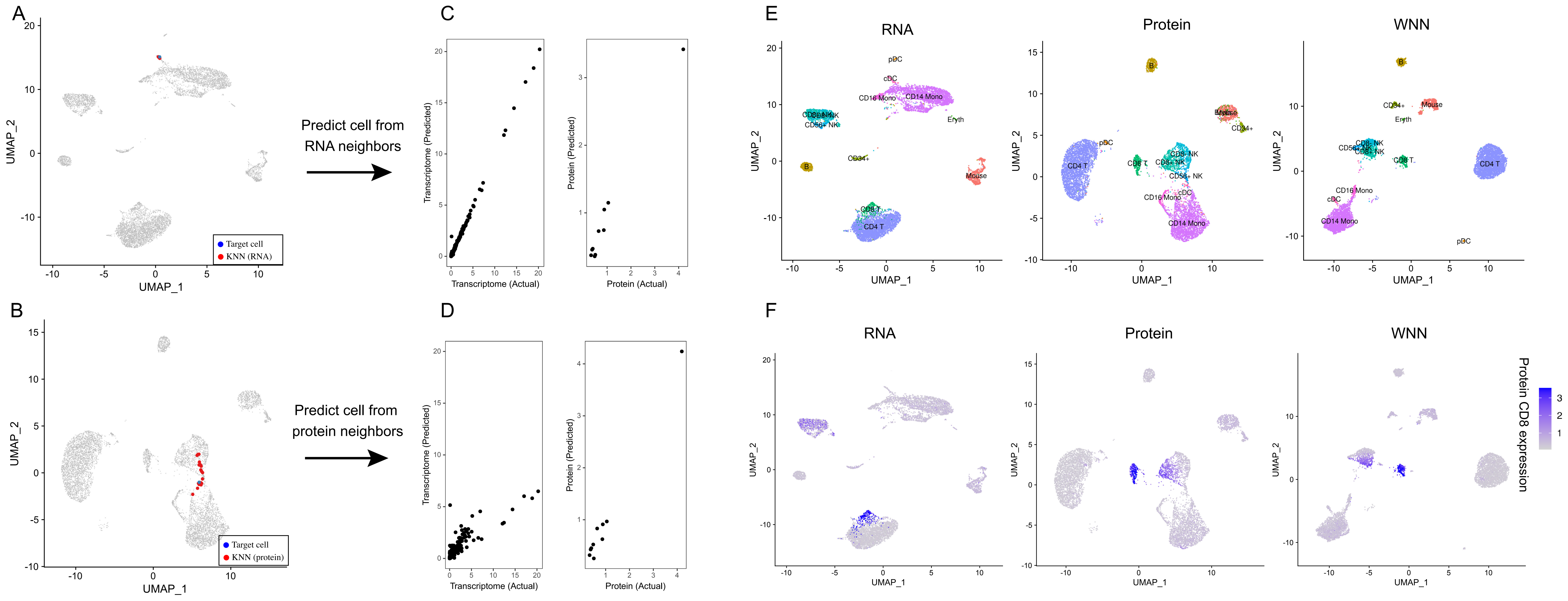


**Supplementary Figure 1: Weighted Nearest Neighbor analysis on a CITE-seq dataset of cord blood mononuclear cells**

**(A, B)** Independent analysis of transcriptome (A) and protein (B) modalities from a CITE-seq analysis of cord blood mononuclear cells. Panels A-D correspond to Figure 1A-D, but the target cell is a dendritic cell instead of a CD8 T cell. Blue dot marks the same target dendritic cell in (A) and (B). Red dots denote the k=20 nearest neighbors to the target dendritic cell based on the transcriptome (A) or protein (B) modalities. **(C)** The RNA neighbors are averaged together to predict the molecular contents of the target dendritic cells. Since the RNA neighbors are all dendritic cells, the predicted values are close to the actual measurements. **(D)** Same as in (C), but averaging protein neighbors. Since protein neighbors are a mixture of cell types, there is substantial error between predicted and measured RNA expression. Thus, the RNA data is more informative for characterizing the state of the target cell, and the cell is assigned an increased RNA modality weight. **(E)** RNA, Protein and WNN UMAP visualization for this dataset. Cells are annotated by their WNN-assigned labels. Visualizations are the same as in Figure 1, but all cell types are labeled on the UMAP plots for greater clarity. **(F)** Feature plot of CD8 protein expression on all three UMAP visualizations, showing that WNN and ADT analyses help to separate CD4 and CD8 T cells, and also identify additional heterogeneity within NK cells.

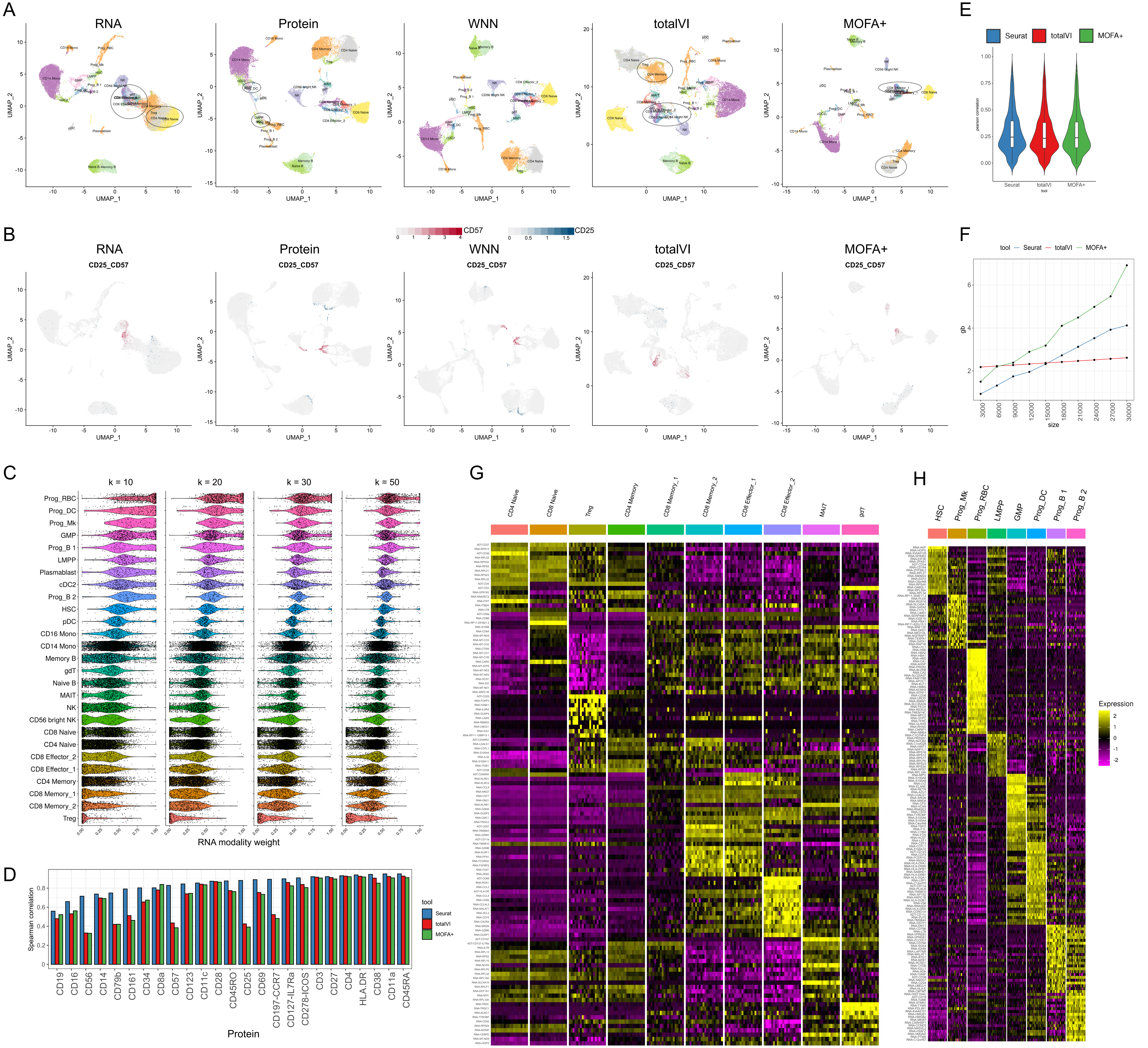

**Supplementary Figure 2: Benchmarking and robustness analysis for WNN integration on a CITE-seq dataset of human bone marrow mononuclear cells (BMNC).**

**(A)** UMAP visualizations of the BMNC dataset based on five analytical strategies: independent RNA analysis, independent Protein analysis, WNN, totalVI and MOFA+. Cell annotations are derived from WNN analysis, which reflect distinct molecular states (see heatmaps in (G-H)). Dashed ovals indicate regions in each analysis where cell states are intermixed. **(B)** Expression of CD25 and CD57 in these five UMAP visualizations. In WNN analysis, cells that are positive for these proteins are correctly determined to be neighbors of each other, and therefore separate in UMAP visualization. **(C)** Robustness analysis for k in the WNN procedure. We varied the number of single-cell RNA modality weights across different number of k-nearest neighbors used (k = 10, 20, 30, 50) on the BMNC dataset, and show single-cell violin plots of the resulting RNA modality weight. We observe only minor fluctuations when varying k within this range. **(D)** Benchmarking WNN against totalVI and MOFA+. The integrated latent space defined by WNN most accurately reconstructs expression levels for all 25 proteins. Same as Figure 2D but showing Spearman correlation instead of Pearson correlation. **(E)** When using the integrated latent space to reconstruct 2000 variable features in the transcriptome, all three methods exhibit equivalent performance. Figure shows boxplot of Pearson correlation between predicted and measured values for 2,000 features. Benchmarking metrics are described further in Supplementary Methods. **(F)** Memory usage for all three methods as a function of the size of the input dataset. **(G)** Heatmap of WNN-annotated T cell states. Features include the best RNA and protein features identified by differential expression. Heatmap displays pseudobulk averages where cells are grouped by cell type, human donor, and technical replicate, and demonstrates that markers are repeatedly detected across samples and replicates. **(H)** Same as in (G) but for progenitor cell states.

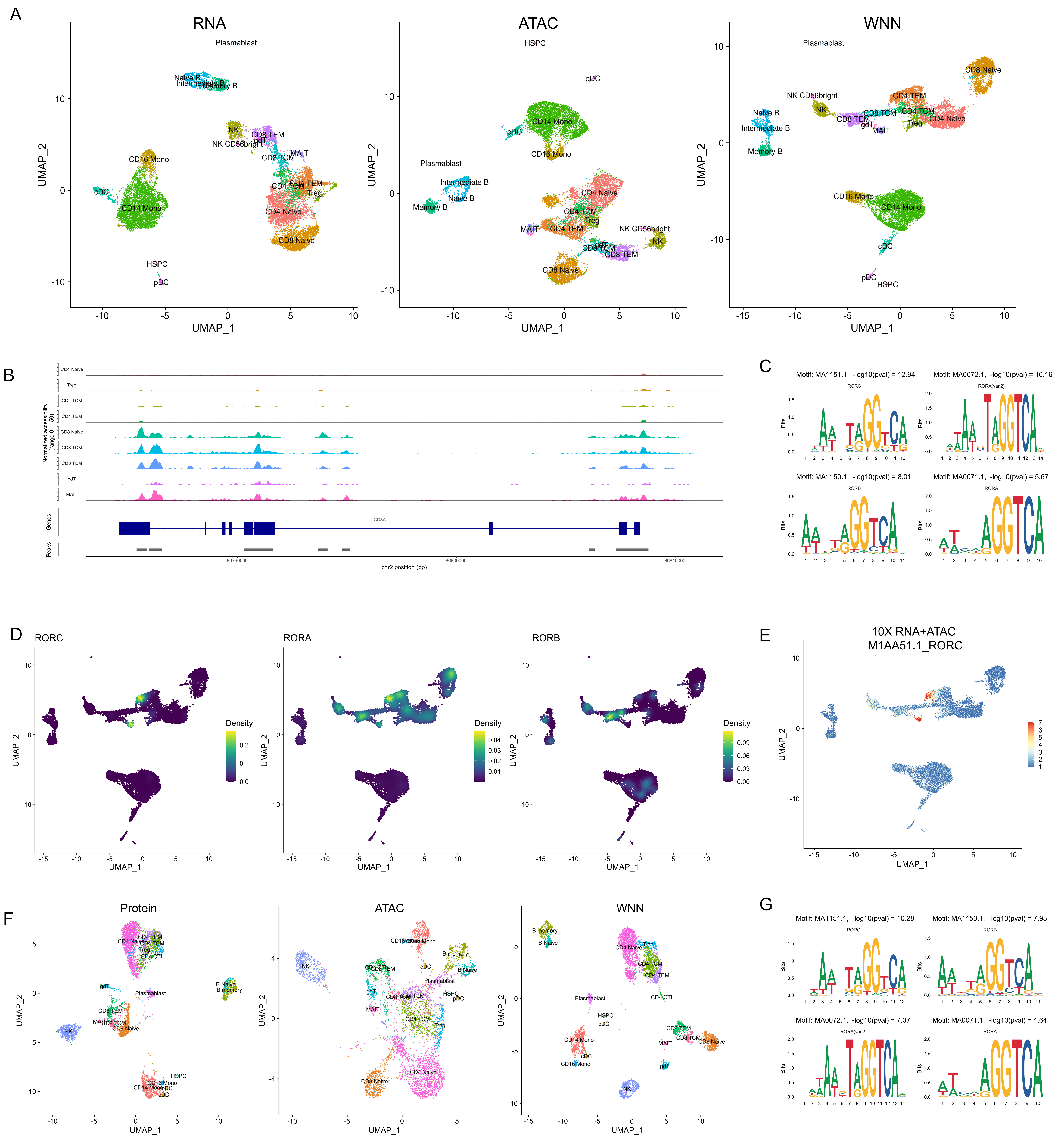


**Supplementary Figure 3. Applying WNN to additional multimodal technologies**

**(A)** Analysis of a publicly available dataset of 11,351 PBMC processed with the 10x Genomics Multiome ATAC+RNA kit. UMAP visualizations of RNA and ATAC-seq data, as well as integrated WNN analysis. Cells are labeled by their WNN-annotated clusters. **(B)** Visualization of pseudobulk chromatin accessibility tracks of the CD8A locus for eight T cell subsets. Multiple peaks clearly separate CD8+ and CD8- T cells, exemplifying the information in ATAC-seq that can enhance parallel RNA measurements for defining cell states. **(C)** Enriched motifs within MAIT-specific open chromatin regions. Since multiple transcription factors (i.e. RORA, RORB, RORC) have very similar binding motifs, each exhibits strong evidence of enrichment. **(D)** Density plots, produced by the Nebulosa package, showing the RNA expression of RORC, RORA and RORB. **(E)** Visualization of RORC motif activity, as calculated by chromVAR, which mirrors the expression of the RORC as shown in (D). **(F)** Analysis of a published ASAP-seq dataset of 4,725 human PBMC where chromatin accessibility and surface expression of 227 surface proteins are simultaneously measured. UMAP visualizations of ATAC and protein data, as well as integrated WNN analysis. Cells are labeled by their WNN-annotated clusters. **(G)** Enriched motifs within MAIT-specific open chromatin regions in the ASAP-seq dataset are concordant with those identified in ATAC+RNA analysis.

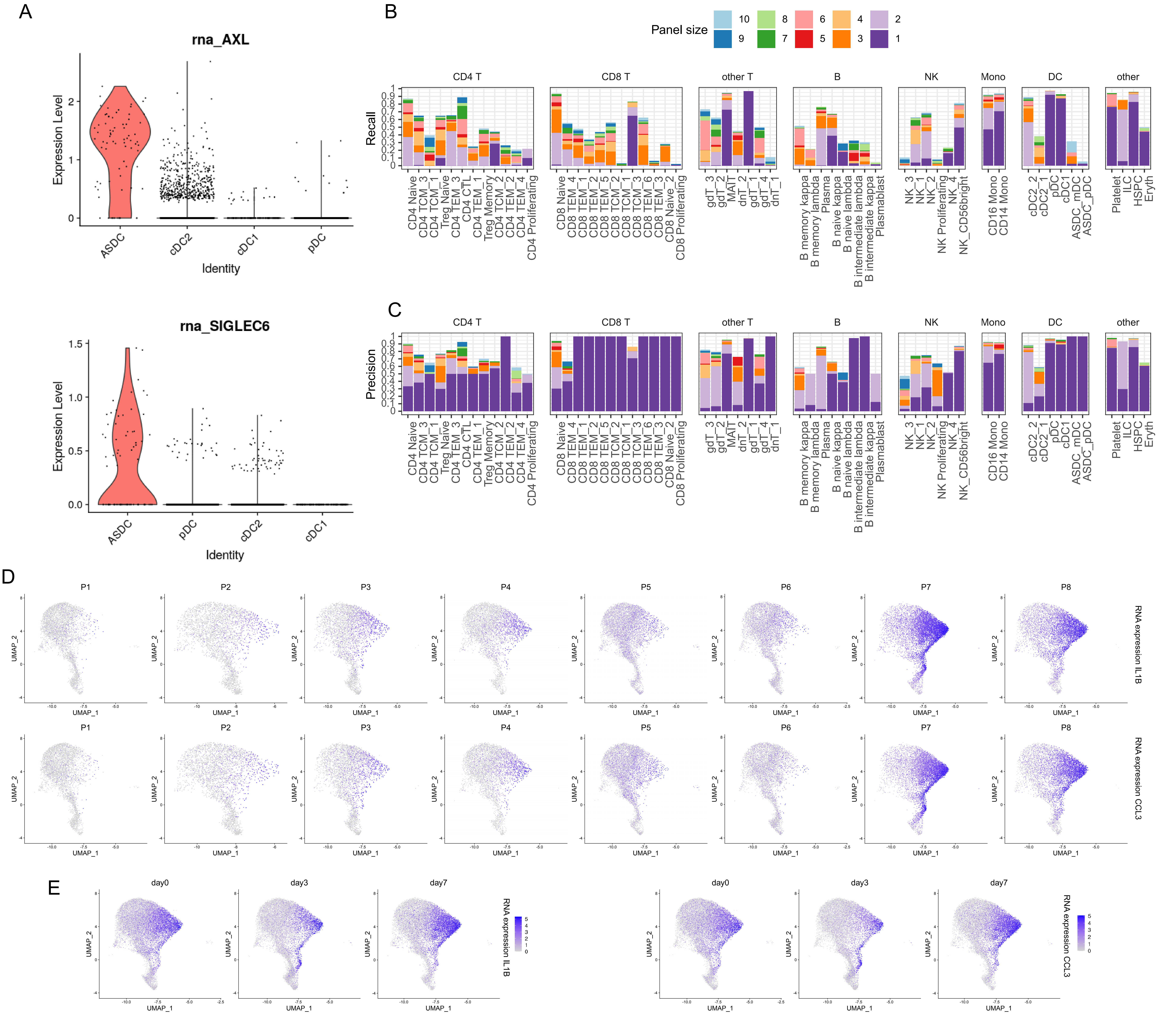


**Supplementary Figure 4: Identifying targeted immunophenotype panels for cell states**

**(A)** RNA expression of two canonical markers of AXL+ SIGLEC6+ dendritic cells (ASDC). Both markers were specifically enriched in the ASDC cells compared to other DC subsets. **(B-C)** For each of the 57 clusters, we computed targeted immunophenotype panels using forward selection coupled with logistic regression. In Figure 4C we visualize the level of enrichment for each cluster based on panels of one to ten markers. Here, we show precision and recall metrics based on logistic regression, using a decision boundary of 0.5. These data demonstrate that while we can achieve substantial enrichment with small panels, isolating pure and homogeneous populations based on small marker panels remains challenging for some clusters. **(D-E)** Additional heterogeneity in the expression of inflammatory genes in monocyte populations. Only CD14+ and CD16 monocytes are shown. Heterogeneous expression of these genes is exhibited in multiple, but not all, volunteers. This heterogeneity was not related to the vaccination timecourse, as shown in (E).

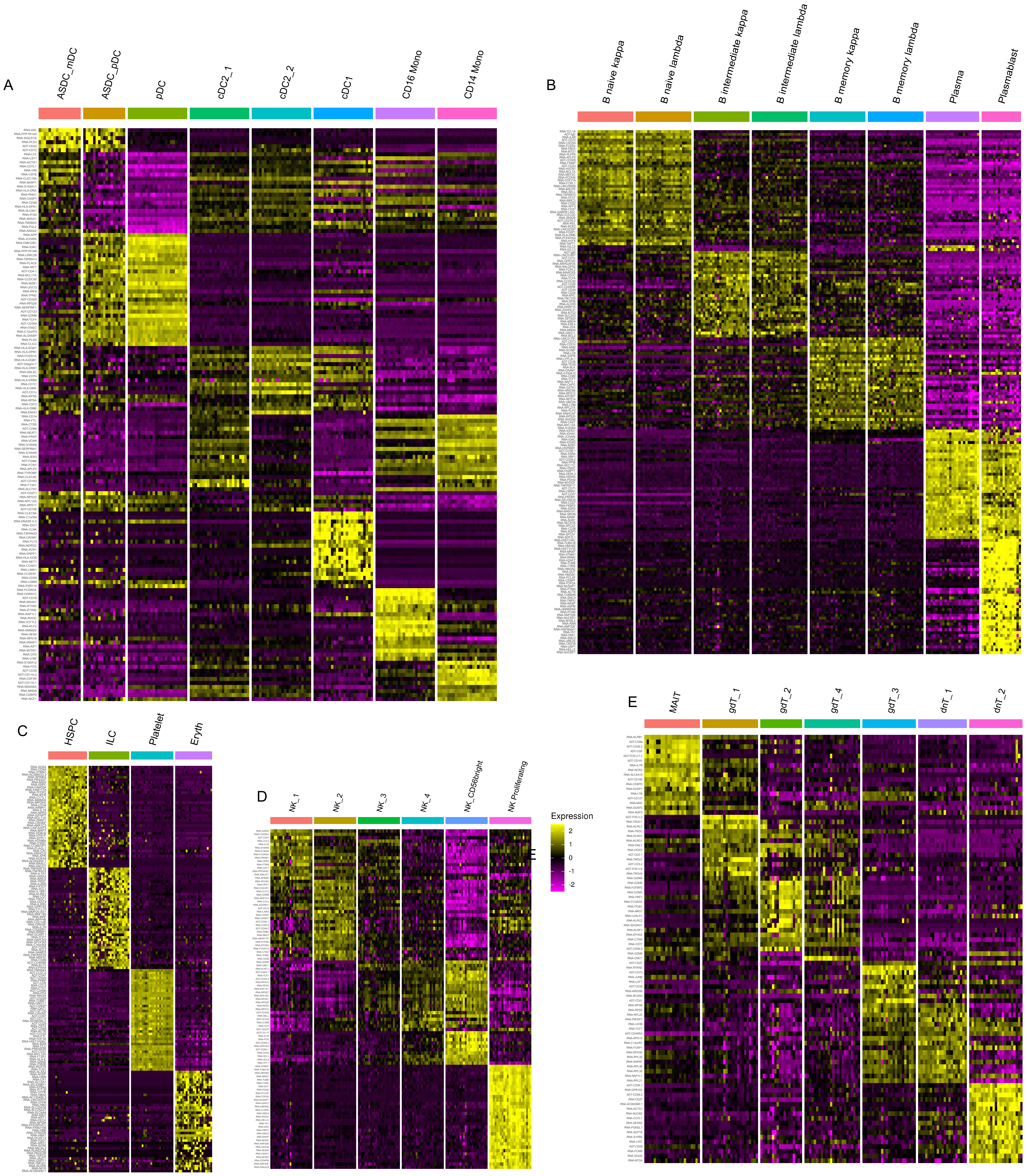


**Supplementary Figure 5. Additional heatmaps for other immune cell states of Multimodal PBMC**

**(A)** Heatmap of myeloid cells states. Features include the best RNA and protein features identified by differential expression. Heatmap displays pseudobulk averages where cells are grouped by cell type, human volunteer, and vaccination time point and demonstrates that markers are repeatedly identified across samples. **(B)** Same as in (A) but for B cell states. B cell states are subdivided by their mutually exclusive expression of kappa or lambda light chain, with distinguishing markers including IGKC, IGLC3, IGLC3. **(C)** Same as in (A) but for unconventional T cells states. **(D)** Same as in (A) but for NK cells states. **(E)** Same as in (A) but for other cells states. Heatmaps for CD4⁺ T cell and CD8⁺ T cell states are shown in Figure 4A-B.

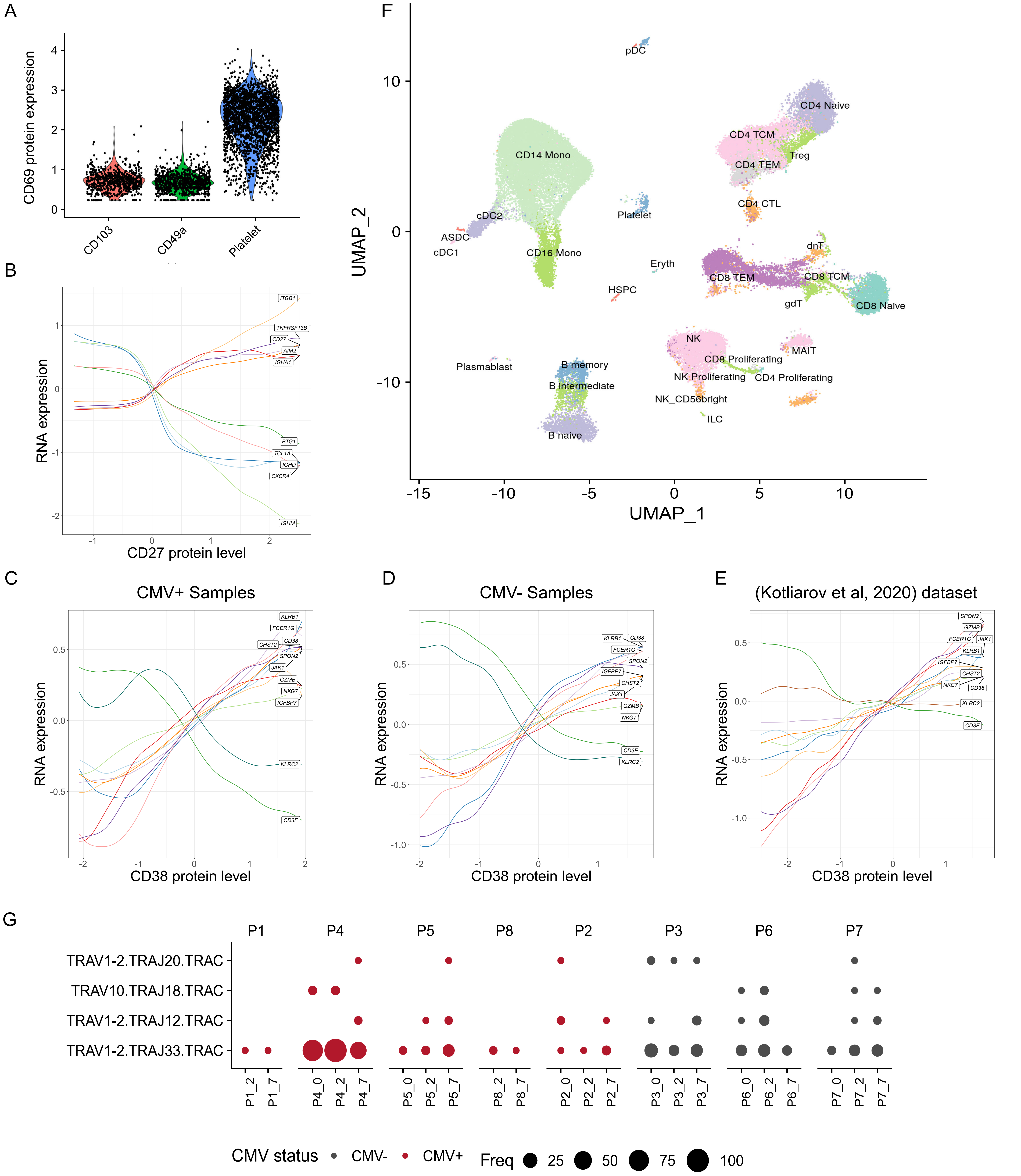

**Supplementary Figure 6. Additional heterogeneity within lymphoid populations**

**(A)** Protein expression of canonical resident lymphocyte marker CD69 in CD8+ CD103+, CD8+ CD49a+ T cells populations. Neither population is positive. Platelets are included as a positive control, as CD69 is constitutively expressed on these cells. **(B)** Naïve, intermediate and memory B cells are ordered by their quantitative level of CD27 protein expression. Rolling averages for the expression of genes that correlate positively or negatively with CD27 are shown as smoothed lines. **(C-E)** Same as Figure 5G, but after splitting the eight volunteers into five CMV+ (C) and three CMV- (D) samples (Supplementary Table 3). We observe concordant trends in both subsets, as well as an independent CITE-seq dataset (Kotliarov et al, Nature Medicine 2020). **(F)** UMAP visualization of CITE-seq dataset of 49,147 PBMC analyzed with the 10X 5' Immune Profiling kit, which also measures immune repertoires. The dataset has been mapped onto the 3'-defined multimodal reference, allowing cells to be visualized in the same UMAP space as the reference, and cells are labeled based on transferred Level 2 annotations. **(G)** Dot plot showing the overrepresentation of TCRα sequences within cells annotated as MAIT. As expected, we detect the canonical MAIT TRAV1-2-TRAJ33 as the most abundant sequence along with reduced usage of TRAJ12 and TRAJ20. We also detect rare populations of invariant NKT cells (defined by the use of TRAV10.TRAJ18). As expected, and in contrast to the clonotypes reported in Figure 5G, these findings are consistent across volunteers, vaccination timepoints, and CMV status.

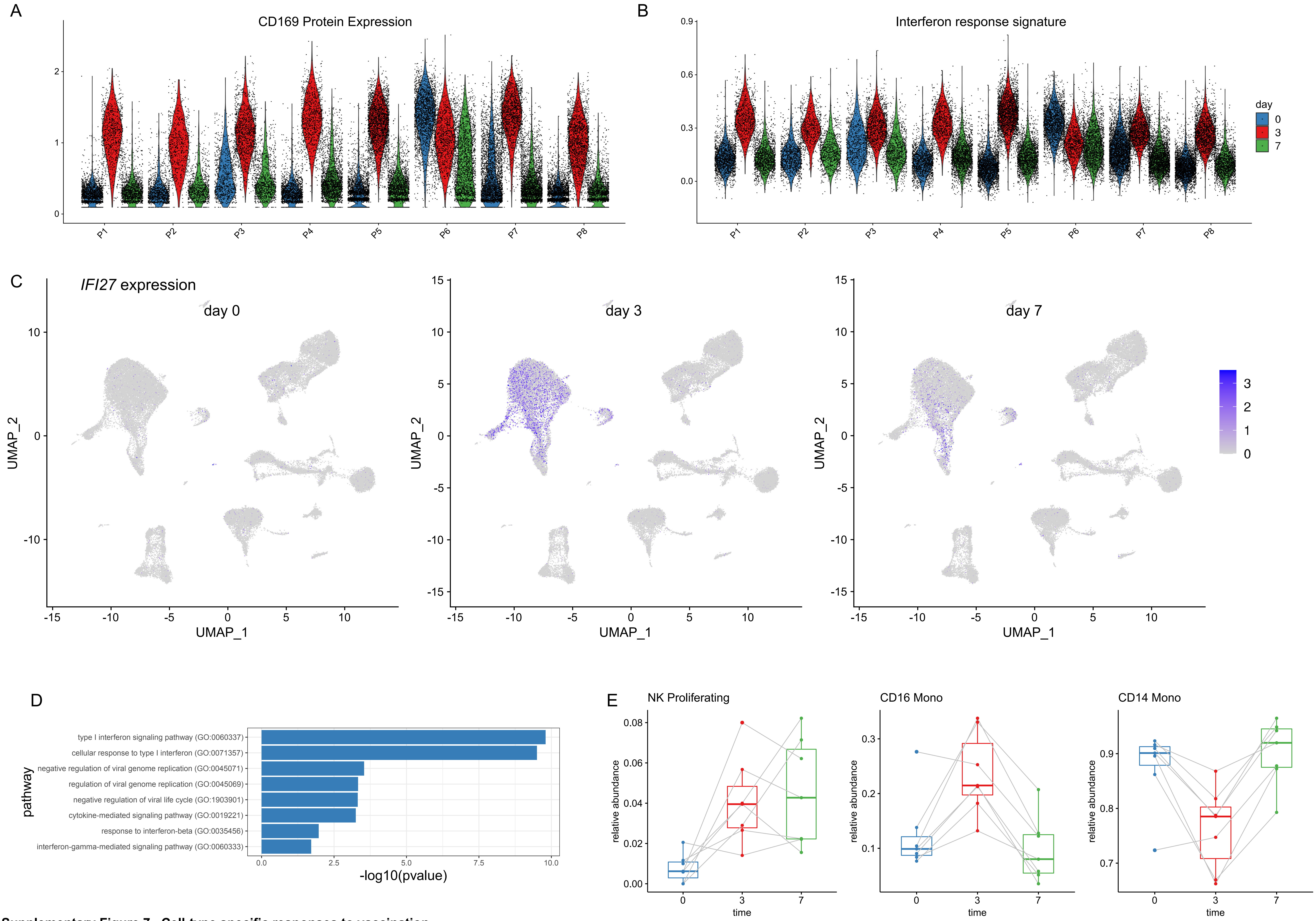


**Supplementary Figure 7. Cell-type specific responses to vaccination**

**(A, B)** Violin plot showing the up-regulation of CD169 protein levels and a module of interferon response genes at day 3. Plot is similar to Figure 6D, but restricted to CD14 Monocytes, and shows the individual response of each volunteer. The response is consistent across all volunteers with one exception (P6), which exhibited signs of a highly activated immune system even prior to vaccination. **(C)** RNA expression of canonical interferon response gene IFI27 across the vaccination time course. The expression of IFI27 increases within particular myeloid populations at day 3, but dampens at day 7. **(D)** Pathway enrichment (enrichR) of the top DE genes between day 0 and day 3 myeloid cells exhibits a clear enrichment for components of the interferon response. **(E)** Same as in Figure 6F, but computed for cells profiled with the 10X 5' kit.

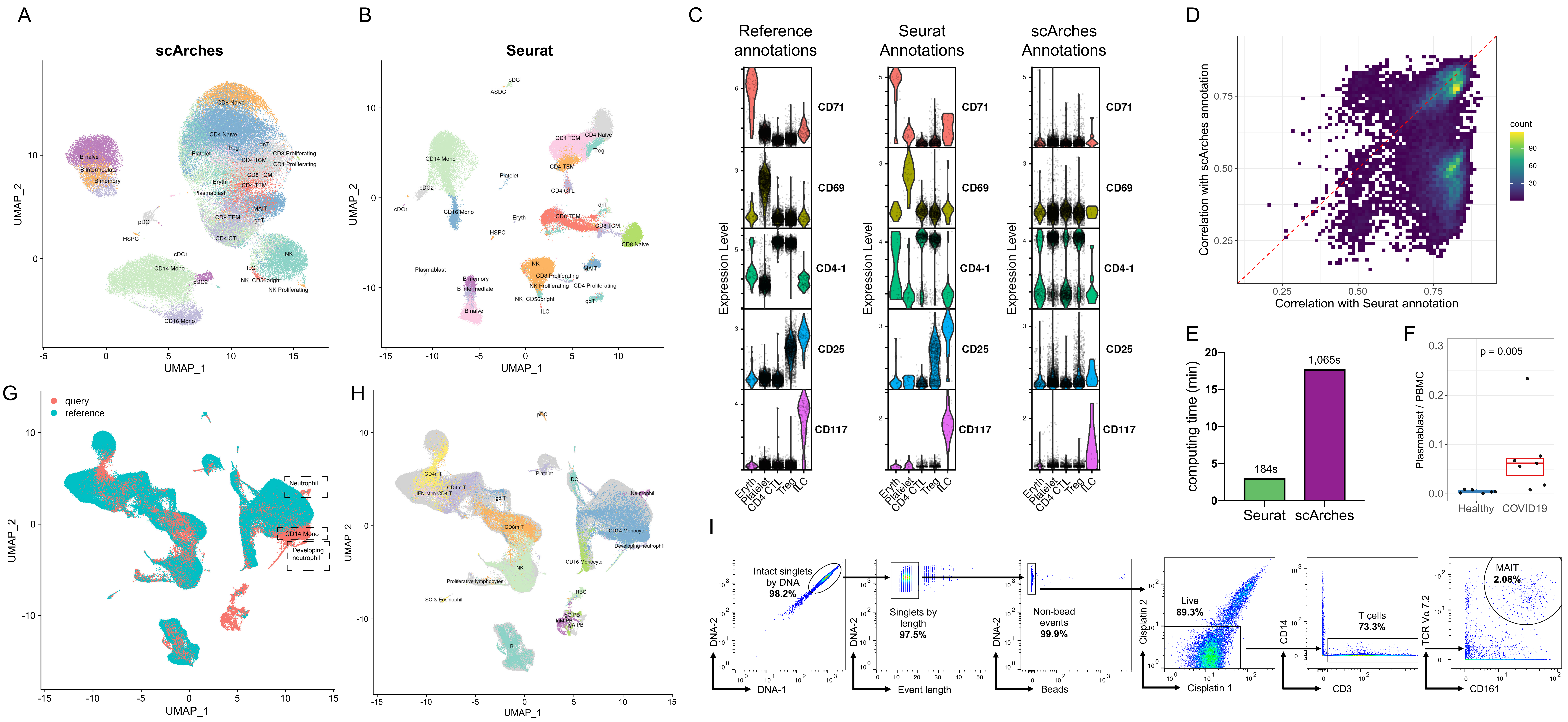


**Supplementary Figure 8. Reference-based mapping of query datasets**

**(A-E)** Benchmarking of Seurat v4 reference-based mapping with scArches. Both methods utilize reference datasets to assist in the interpretation of query data. **(A-B)** UMAP visualizations of reference-based mapping of a human PBMC CITE-seq dataset from (Kotliarov et al, 2020). Cells are label by the annotation that was transferred using each method. The protein data was withheld from mapping and can be used to assess accuracy. **(C)** For five cell types where we observed a high rate of discordant predictions between Seurat and scArches, we visualize the protein expression of key markers in the reference dataset (left), Seurat-transferred annotations (middle), and scArches-transferred annotations (right). In each case, the Seurat annotations provide the most concordant results. For example, cells annotated by Seurat as Treg express CD25 protein, while cells annotated by scArches as Treg do not. **(D)** For all 17,480 (32.9%) of query cells where Seurat and scArches returned different annotations based on the transcriptome, we calculated protein-based classification metrics to determine the support for each result (Supplementary Methods). In 73.8% of cases, we observe stronger support for the Seurat annotation. **(E)** Computing time for reference-mapping of (Kotliarov et al, 2020) onto the multimodal reference. **(F)** The abundance of plamablasts increases during COVID-19 response. p-value is computed using an unpaired Wilcoxon test. Annotations were derived from reference-based mapping, and confirm the result reported in (Wilk et al, 2020). **(G)** 'de novo' UMAP (Supplementary Methods) visualization of the dataset from (Wilk et al., 2020) after reference-mapping. Concordant cell types are identified between query and reference data with three exceptions, denoted with dashed rectangles. **(G)** Same as in (E), but cells are colored by their unsupervised label as described in (Wilk et al., 2020). These results demonstrate that developing and differentiated neutrophils, which are not present in the reference, remain distinct after reference-based mapping. Additionally, a population of CD14+ Monocytes that has severe transcriptional responses to COVID-19 is also highlighted in this analysis. **(I)** Gating strategy used to identify MAIT cells in mass cytometry experiments.
