## Supplementary Methods for "Integrated analysis of multimodal single-cell data"

### HIV vaccine trial specimens

HVTN 087 (NCT01578889) was a phase 1a HIV vaccine trial that tested intramuscular electroporation of a DNA vaccine with or without IL-12 adjuvant delivered as a plasmid at months 0, 1 and 3 followed by boosting with VSV-vectored HIV *gag* vaccine at month 6 [1,2]. Eight participants in this trial were selected for single cell analysis from Group 1 (no IL-12) and Group 3 (1000 mcg IL-12) based on sample availability. Participant demographics and group assignments are listed in Supplementary Table 3. Blood was collected immediately before the first VSV-Gag administration and at sequential timepoints afterwards. PBMC were isolated and cryopreserved as previously described [3].

### Antibody titration, staining, and cleanup

For CITE-seq / TotalSeq-A / 3P scRNA-seq experiments, we pooled together 228 TotalSeq-A antibodies from BioLegend (Supplementary Table 1). In preliminary experiments designed to test the balance of markers in the panel, reads corresponding to 12 antibodies each took up more than 2% of the total sequencing space, and combined together, accounted for half of the total antibody reads. The signal for each of these markers was reduced by addition of a proportional amount of unlabeled antibodies. We recommend the addition of unlabeled antibodies as an effective strategy to modify existing panels to be more robust to sequencing saturation from highly expressed protein markers. CITE-seq antibodies and unlabeled blocking antibodies were then combined in PBS and concentrated with a 50 kDa Amicon filter as per manufacturer's instructions. Post elution, BSA was added to a final concentration of 2%.

For ECCITE-seq / TotalSeq-C / 5P scRNA-seq experiments, we used a combination of antibody:oligo conjugates designed for ECCITE-seq [4], conjugated as described [5], and commercially available TotalSeq-C reagents. 52 ECCITE-seq antibodies [4] were pooled together at a concentration of 1 $\mu$g each per test, and combined with TotalSeq-C reagents for CD45RA and CD45RO at 0.25 $\mu$g each per test. Antibodies were pooled together and concentrated in a 50 kDa Amicon filter as per manufacturer's instructions in PBS. Post elution, BSA was added to a final concentration of 2%.

### CITE-seq staining and sample preparation

To minimize batch effects, for each experiment we processed frozen PBMCs from four different patients at 3 different timepoints (day 0, day 3, and day 7). After thawing, cells were incubated with FcX block (BioLegend) for ten minutes. Cells were then divided into separate aliquots and processed independently for the 3P and 5P protocols.

For the 3P CITE-seq staining protocol, samples were stained simultaneously with the antibody/block pool and a unique hashtag for 30 minutes. Cells were then washed 3 times in staining buffer (2% BSA, 0.01% Tween in PBS) and filtered using a 40 $\mu$m Flowmi filter in PBS and pooled in equal proportions. Cells were loaded into 8 lanes of a 10x Genomics Chip B, at 45,000 cells per lane using the 10x Genomics 3' v3 GEM kit.

For the 5P ECCITE-seq staining protocol, each sample of cells was first stained with a unique hashtag for 30 minutes. Cells were then washed 3 times in staining buffer, pooled together and stained with the antibody panel for 30 minutes. The pool of cells was then washed 3 times in staining buffer and filtered using a 40$\mu$m Flowmi filter in PBS. Cells were loaded into 2 lanes of a 10x Genomics Chip A, at 45000 cells per lane, using the 10x Genomics V(D)J kit (v1).

For both 3P and 5P experiments, first strand cDNA was generated by incubating the emulsions according to the respective 10x Genomics protocol. Emulsions were then broken and nucleic acids recovered Subsequent library preparation steps are detailed in the section below.

*Library prep*

CITE-seq / 3P scRNA-seq:
The 10x 3P v3 protocol was followed according to manufacturer's instructions for cDNA amplification, with the following modifications:

- During cDNA amplification, 0.2 $\mu$M of ADT additive primer (**5'CCTTGGCACCCGAGAATTCC**) and 0.2 $\mu$M of HTO additive primer (5'GTGACTGGAGTTCAGACGTGTGCTC) were added to the reaction.
- During cDNA cleanup, the supernatant from the 0.6x SPRI cleanup was saved and purified with two rounds of 2x SPRI. The eluate was split and used as template for production of ADT and Hashtag libraries:
- Hashtag libraries were generated by PCR using Kapa Hifi Master Mix, 10 $\mu$M 10x Genomics SI-PCR primer (5'AATGATACGGCGACCACCGAGATCTACACTCTTTCCCTACACGACGCTC), and 10 $\mu$M Illumina TruSeq DNA D7xx primer (5'CAAGCAGAAGACGGCATACGAGATxxxxxxxxGTGACTGGAGTTCAGACGTGTGC). Following amplification, Hashtag libraries were and cleaned up with 1.6x SPRI.

Antibody tag libraries were generated by PCR using Kapa Hifi Master Mix, 10 $\mu$M 10x Genomics SI-PCR primer, and 10 $\mu$M TruSeq Small RNA RPIx primer (5'CAAGCAGAAGACGGCATACGAGxxxxxxxxGTGACTGGAGTTCCTTGGCACCCGAGAATTCCA) Following amplification, Antibody tag libraries were and cleaned up with 1.6x SPRI.

*ECCITE-seq / 5P scRNA-seq / immune receptor:*
The 10x Immune Profiling v1 protocol was followed according to manufacturer's instructions for cDNA amplification, with the following modifications:

- During cDNA amplification, 0.2 $\mu$M each of ADT (**5'CCTTGGCACCCGAGAATT*C*C**), HTO (**5'GTGACTGGAGTTCAGACGTGTGC*T*C**), and TotalSeq-C additives (5' CTCGTGGGCTCGGAGATGTGTATAAGAGACAG) were added to the reaction.
- Post cDNA cleanup, a 0.6x SPRI cleanup was performed, where larger cDNA fragments were kept on the beads, and the smaller tag libraries were retained in the supernatant. From the material retained on the beads, a portion of the eluted material was used to generate TCR α/β libraries (as written in the 10x protocol), BCR libraries (as written in the 10x protocol) and TCR γ/δ libraries (as written in the 10x protocol for TCR α/β, with these modifications):
    - 5 $\mu$L of cDNA was taken into the initial reaction
    - For the first PCR, instead of the TCR1 primer mix provided by 10x genomics, we substituted our own mix consisting of primers 5'AGCTTGACAGCATTGTACTTCC and 5'TGTGTCGTTAGTCTTCATGGTGTTCC

    - For the second PCR, instead of the TCR2 primer mix provided by 10x Genomics, we substituted our own of primers consisting of 5'TCCTTCACCAGACAAGCGAC and 5'GATCCCAGAATCGTGTTGCTC
- The 0.6X SPRI supernatant remaining following cDNA cleanup was subjected to 2 rounds of 2x SPRI. The eluate was split into three reactions for tag library production:
    - Hashtag libraries were created by performing a PCR reaction consisting of Kapa Hifi Master Mix, 10 $\mu$M 10x Genomics SI-PCR primer, and 10 $\mu$M Illumina TruSeq DNA D7xx primer.
    - Antibody libraries (for homemade conjugates) were created by performing a PCR reaction with Kapa Hifi Master Mix, 10 $\mu$M 10x Genomics SI-PCR primer, and 10 $\mu$M TruSeq Small RNA RPIx primer.
    - TotlaSeq-C antibody libraries were created by performing a PCR reaction with 2x Kapa Hifi Master Mix, 10 $\mu$M 10x Genomics SI-PCR primer, and 10 $\mu$M Nextera indexing primer (CAAGCAGAAGACGGCATACGAGATxxxxxxxxGTCTCGTGGGCTCGGAGATGTGTATAAGAGACAG).
    -

*Sequencing*

For 3P libraries, the samples were pooled in a ratio of 80% RNA, 12% ADT, and 8% HTO.
For the 5P libraries, the samples were pooled in a ratio of 70% RNA, 12% ADT, 8% HTO, 5% of TCR libraries (with equal amounts of α/β and γ/δ libraries), and 5% of BCR libraries. 3P and 5P libraries were then pooled together in equal amounts and sequenced on an Illumina Novaseq S4 flowcell.

**Validation of targeted immunophenotype panels experiments**

Commercially available cryopreserved PBMCs (AllCells) were thawed into DMEM with 10% FBS. Two million cells per condition (4 conditions) were spun down in Eppendorf tubes at 4 ºC for 5 min at 400 *g*, and resuspended in 100 $\mu$l PBS with 2% BSA. Each aliquot was incubated for 10 minutes with 10 $\mu$L of FcX block, followed by staining with flow cytometry antibodies (BioLegend) on ice for 30 minutes. Cells were washed three times with PBS with 2% BSA. Samples were then gated as described below and sorted directly into Buffer RLT (Qiagen).

Antibodies used (all at 5*u*L per condition unless otherwise noted):

| Specificity | Fluorophore | Clone | Note |
|---|---|---|---|
| CD3 | AF488 | UCHT1 | 10*u*L of antibody used |
| CD8 | APC-Cy7 | SK1 | |
| CD4 | AF700 | RPA-T4 | |
| CD57 | PE | QA17A04 | |
| CD56 | APC | 5.1H11 | |
| CD103 | BV421 | Ber-ACT8 | |
| Integrin 7 | PE | FIB504 | |
| CD49a | APC | TS2/7 | |

| | | |
|---|---|---|
| CD43 | PE | CD43-10G7 |

Gating conditions for each of the validation experiments are shown in Figure 4D, E.

Post sorting, samples were each split into quintuplicates, and then cleaned up with 2x SPRI. Samples were then brought into reverse transcription in an adaptation of SMARTseq2 [6] and SCRB-seq [7] as described here: dx.doi.org/10.17504/protocols.io.nkgdctw

The pooled library was sequenced on an Illumina Nextseq (50 R1, 8 index, 34 R2). Post base calling, samples were aligned using a wrapper for DropSeqTools against the human reference hg19 to generate RNA counts matrices.

To assess the agreement between single-cell datasets and bulk-sorted experiments, we examined the top DE genes separating our gated populations in the CITE-seq reference dataset. We next visualized the relative expression of these genes in the heatmaps in Figure 4D, E. The bulk-sorted populations exhibited highly concordant relative expression patterns for DE genes as we observed in CITE-seq data.

**Flow cytometry analysis of whole blood**

Whole blood collected immediately before the first VSV *gag* administration and then 1 and 7 days after was stained in TruCOUNT tubes as previously described [1, 8, 9] using the following antibody staining panel: (antibodies from BD Biosciences, unless otherwise indicated): CD14–V450, CD19–V450, CD45–AmCyan, CD4–FITC, CD8–PerCP-Cy5.5, CD123–PE, HLA-DR–ECD (Beckman Coulter), CD86–PE-Cy5, CD56–PE-Cy7, CD11c–APC, CD3–Alexa700 and CD16–APC-Cy7. We used these measurements in Figure 6G-H to validate changes in cell type abundance that were detected by scRNA-seq.

**Determination of cellular responses to CMV**

Intracellular cytokine staining assays were conducted as described in [2] and the proportion of $CD8^+$ T cells expressing IL-2 and/or IFN-$\gamma$ after stimulation with a CMV peptide pool as well as the response call are listed are in Supplementary Table 3**.**

**Mass cytometry**

PBMC from patients with nasopharyngeal swab PCR-confirmed COVID-19 and healthy controls were thawed into warm RPMI (HyClone/Thermo Scientific) supplemented with 10% FBS and 0.5 x 106 cells per sample were transferred into a 96-well plate for staining. Cells were stained as previously described in [10], using the panel described in [11] with the addition of the following antibodies to identify MAIT cells: anti-CD161 (DX12, BD Biosciences) conjugated on 141Pr and TCR V$\alpha$7.2 (clone 3C10, Biolegend) conjugated on 162Dy. Antibodies were conjugated using MaxPar® X8 Conjugation Kits (Fluidigm, South San Francisco, CA, USA) or purchased pre-conjugated from Fluidigm. With the exception of the antibodies added, the immune profiling panel was premixed and frozen at -80°C in order to ensure antibody stability and minimize differential staining between batches as described in [11]. Briefly, cells were washed with PBS (Rockwell) and resuspended in 25 mM cisplatin (Enzo, Farmingdale, NY, USA) for sixty seconds to stain for viability before being quenched with undiluted FBS. Samples were multiplexed by staining with CD45-Pd barcodes as previously described [12], washed thoroughly in CyFACS buffer (PBS, 0.1% BSA, 2mM EDTA, 0.05% sodium azide), and pooled into sets of barcodes. Barcoded samples were then stained with all antibodies for 30 minutes, washed with CyFACS buffer, and fixed in 2% Paraformaldehyde

(Electron Microscopy Sciences, Hatfield, PA, USA) for 20 minutes at room temperature. Fixed cells were permeabilized with 1x eBiosciences Permeabilization Buffer. Manufacturer ThermoFisher Scientific (Waltham, MA). Samples were washed, resuspended in 2% PFA containing iridium intercalator (Fluidigm), and stored at 4°C until acquisition (within 3 days of staining). On the day of acquisition, samples were washed once with PBS and thrice with Milli-Q water before being resuspended in 1× EQ Beads (Fluidigm) and collected on a Helios mass cytometer (Fluidigm).

Prior to analysis, fcs files were debarcoded and bead-normalized with EQ beads using the Premessa package in the open-source statistical software R as previously described [13]. FlowJo v10.7.1 was used to visualize the data and used to gate out beads, dead cells, doublets, and cell debris. MAIT cells were identified by expression of CD3, CD161, and TCR Vα 7.2 as shown in Supplementary Figure 8.

**Weighted Nearest Neighbor Analysis**

The weighted nearest neighbor (WNN) procedure implemented in Seurat v4 is designed to integrate multiple types of data that are collected in the same cells to define a single unified representation of single-cell multimodal data. For each cell, the procedure learns a set of modality weights, which reflect the relative information content for each data type in that cell. This enables the generation of a WNN graph: for each cell, this graph denotes the most similar cells in the dataset based on a weighted combination of protein and RNA similarities. The WNN graph can be used as input for common downstream analytical tasks including tSNE or UMAP visualization, graph-based clustering, and the identification of developmental trajectories.

Our approach consists of four broad steps, as explained in detail below: (1) Constructing independent *k*-nearest neighbor (KNN) graphs for both modalities. (2) Performing within and across-modality prediction (3) Calculating cell-specific modality weights. (4) Calculating a WNN graph.

All methods are implemented in our open-source R package Seurat (www.satijalab.org/seurat, www.github.com/satijalab/seurat)

*Constructing k-nearest neighbor graphs for each modality*

The WNN procedure begins by first applying standard analytical workflows to each modality independently and constructing KNN graphs for each one. In this manuscript we analyze data falling into three categories: measurements of single-cell gene expression, single-cell surface protein expression, and single-cell chromatin accessibility (ATAC-seq). For most analyses in this manuscript, we use a default value of $k$=20, which is also the default value of $k$ in the standard Seurat clustering workflow. For the analysis of the multimodal PBMC atlas, due to the substantial size of the dataset, we used a value of $k$=30. In Supplementary Figure 2, we show that we obtain very similar results from the WNN procedure when varying $k$ across a series of values ranging from 10 to 50.

For clarity, we overview the analytical workflows for each data type below:

Single-cell gene expression: We analyze scRNA-seq data using standard pipelines in Seurat which include normalization, feature selection, and dimensional reduction with PCA. We then construct a KNN graph after dimensional reduction.

We emphasize that WNN analysis can leverage any scRNA-seq preprocessing workflow that generates a KNN graph. For example, users can preprocess their scRNA-seq data with a variety of normalization tools including log-normalization, scran [14] or SCTransform [15], and can utilize alternative dimensional reduction procedures such as factor analysis or variational autoencoders. In this manuscript, we use workflows that are available in the Seurat package, and detail parameter settings for each analysis later in this document.

Single-cell cell surface protein level expression: We analyze single-cell protein data (representing the quantification of antibody-derived tags (ADTs) in CITE-seq or ASAP-seq data) using a similar workflow to scRNA-seq. We normalize protein expression levels within a cell using the centered-log ratio (CLR) transform, followed by dimensional reduction with PCA, and subsequently construct a KNN graph. Unless otherwise specified, we do not perform feature selection on protein data, and use all measured proteins during dimensional reduction.

Single-cell chromatin accessibility: We analyze single-cell ATAC-seq data using our previously described workflow [16], as implemented in the Signac package. We reduced the dimensionality of the scATAC-seq data by performing latent semantic indexing (LSI) on the scATAC-seq peak matrix, as suggested by Cusanovich and Hill et al. [17]. We first computed the term frequency-inverse document frequency (TF-IDF) of the peak matrix by dividing the accessibility of each peak in each cell by the total accessibility in the cell (the "term frequency"), and multiplied this by the inverse accessibility of the peak in the cell population. This step 'upweights' the contribution of highly variable peaks and downweights peaks that are accessible in all cells. We then multiplied these values by 10,000 and log-transformed this TF-IDF matrix, adding a pseudocount of 1 to avoid computing the log of 0. We decomposed the TF-IDF matrix via SVD to return LSI components, and scaled LSI loadings for each LSI component to mean 0 and standard deviation 1.

As described for scRNA-seq analysis, while we use Seurat and Signac functions in this manuscript, any analytical workflow that computes a KNN graph for surface protein or chromatin accessibility data can also be used in the first step of WNN analysis.

*Performing within and across-modality predictions*

Suppose we have a CITE-seq dataset where two modalities, RNA and protein, are measured in each single cell. From the previous step, we define the following:

$r_i$: L2-normalized low-dimensional vector representing the RNA profile for cell $i$

$p_i$: L2-normalized low-dimensional vector representing the protein profile for cell $i$

$\{knn_{r,i,1} \dots knn_{r,i,k}\}$ : the set of k-nearest RNA neighbors for cell $i$

$\{knn_{p,i,1} \dots knn_{p,i,k}\}$: the set of k-nearest protein neighbors for cell $i$

We average the low-dimensional profiles of each neighbor set, which represents a prediction for the molecular contents for cell $i$ based on its local neighborhoods. We perform both within-modality and across-modality prediction:

Within-modality prediction:

$\hat{r}_{i,knn_r} = \frac{\sum_{j=1}^{k} r_{knn_{r,i,j}}}{k}$ : prediction of RNA profile for cell $i$, based on RNA neighbors

$$\hat{p}_{i,knn_p} = \frac{\sum_{j=1}^{k} p_{knn_{p,i,j}}}{k}$$ : prediction of protein profile for cell $i$, based on protein neighbors

Cross-modality prediction:

$$\hat{r}_{i,knn_p} = \frac{\sum_{j=1}^{k} r_{knn_{p,i,j}}}{k}$$ : prediction of RNA profile for cell $i$, based on protein neighbors

$$\hat{p}_{i,knn_r} = \frac{\sum_{j=1}^{k} p_{knn_{r,i,j}}}{k}$$ : prediction of protein profile for cell $i$, based on RNA neighbors

_Calculating cell-specific modality weights_

We next calculate the similarity between predicted values for each cell $\hat{r}_i$ and $\hat{p}_i$, and the actual values $r_i$ and $p_i$. We first compute Euclidean distances between predicted and actual values, and next convert these to affinities using the exponential kernel defined in [18].

$$\theta_{rna}(r_i, \hat{r}_{i,knn_r}) = \exp\left( \frac{-\max(d(r_i, \hat{r}_{i,knn_r}) - d(r_i, r_{knn_{r,i,1}}), 0)}{\sigma_{r,i} - d(r_i, r_{knn_{r,i,1}})} \right)$$

affinity between $r_i$ and predicted RNA profile (based on RNA knn)

$$\theta_{rna}(r_i, \hat{r}_{i,knn_p}) = \exp\left( \frac{-\max(d(r_i, \hat{r}_{i,knn_p}) - d(r_i, r_{knn_{r,i,1}}), 0)}{\sigma_{r,i} - d(r_i, r_{knn_{r,i,1}})} \right)$$

affinity between $r_i$ and predicted RNA profile (based on protein knn)

$$\theta_{protein}(p_i, \hat{p}_{i,knn_p}) = \exp\left( \frac{-\max(d(p_i, \hat{p}_{i,knn_p}) - d(p_i, p_{knn_{p,i,1}}), 0)}{\sigma_{p,i} - d(p_i, p_{knn_{p,i,1}})} \right)$$

affinity between $p_i$ and predicted protein profile (based on protein knn)

$$\theta_{protein}(p_i, \hat{p}_{i,knn_r}) = \exp\left( \frac{-\max(d(p_i, \hat{p}_{i,knn_r}) - d(p_i, p_{knn_{p,i,1}}), 0)}{\sigma_{p,i} - d(p_i, p_{knn_{p,i,1}})} \right)$$

affinity between $p_i$ and predicted protein profile (based on RNA knn)

In the equations above, $d$ represents the Euclidean distance metric, and $\sigma_{r,i}$ and $\sigma_{p,i}$ represent the bandwidth of the RNA and protein kernels for cell $i$. A commonly used approach is to set the bandwidth of a kernel to reflect the distance between a cell and its $k$-th nearest neighbor, resulting in an adaptive bandwidth that is specific to each cell [19][20]. However, the value of $k$ used to compute this bandwidth is typically fixed across all cells. We considered that cells originating from rare states should not have the same bandwidth constraint as cells originating from abundant states, and therefore considered a modified approach to select kernel bandwidths.

Our approach is inspired by the concept of large margin nearest neighbors, which aims to identify kernel bandwidths that separate data points in the same class from those in different classes, even if the classes are closely related [21]. In the context of unsupervised single-cell analysis (where the data points are unlabeled), we aim to identify a kernel bandwidth that groups together cells in the same state, yet divides cells that originate from closely related (but different) states.

Recent work has clearly demonstrated that KNN-graphs are prone to the formation of spurious edges, which represent links between cells that share some similarity molecular profiles, but are not in a matched molecular state [22]. However, it is possible to identify these spurious edges through the use of the Jaccard metric. This identifies the number of shared nearest neighbors between two cells, thereby exploiting the local density of each data point to separate well-supported from spurious edges.

For each cell $i$, we therefore aim to identify the 20 cells in the dataset with the *lowest* non-zero Jaccard similarity. We expect that these represent cells that exhibit some similarity with cell $i$, but are unlikely to reside in the same molecular state. If more than 20 cells share the same Jaccard value, we select the 20 with the furthest euclidean distance to cell $i$. We take the average of the Euclidean distances from cell $i$ to the 20 selected cells, and set this as the cell-specific kernel bandwidth.

*Calculating cell-specific modality weights*

We next calculate the ratio between the affinities for $r_i$ with predictions based on RNA neighbors, and predictions based on protein neighbors. A large ratio suggests that the local neighborhood of the cell, as defined by its RNA neighbors, better reflects its molecular state. We calculate the analogous ratio for protein affinities for $p_i$. In both cases, we add a small $\varepsilon$ ($10^{-4}$) to the denominator to avoid numerical errors.

$$s_{rna}(i) = \frac{\theta_{rna}(r_i, \hat{r}_{i,knn_r})}{\theta_{rna}(r_i, \hat{r}_{i,knn_p}) + \varepsilon}, \quad s_{protein}(i) = \frac{\theta_{protein}(p_i, \hat{p}_{i,knn_p})}{\theta_{protein}(p_i, \hat{p}_{i,knn_r}) + \varepsilon}$$

Finally, we normalize these values with a softmax transformation. The resulting two values are non-negative, and together sum to 1. We refer to these as cell-specific modality weights.

$$w_{rna}(i) = \frac{e^{s_{rna}(i)}}{e^{s_{rna}(i)} + e^{s_{protein}(i)}}, \quad w_{protein}(i) = \frac{e^{s_{protein}(i)}}{e^{s_{rna}(i)} + e^{s_{protein}(i)}}$$

*Calculating a WNN graph*

We leverage the cell-specific modality weights calculated above to define a new similarity metric between any two cells, which reflects a weighted combination of RNA and protein affinities. For two cells $i$ and cell $j$, we define their weighted similarity as:

$$\theta_{weighted}(i,j) = w_{rna}(i)\theta_{rna}(r_i, r_j) + w_{protein}(i)\theta_{protein}(p_i, p_j)$$

We then construct a WNN graph, defined as a KNN graph constructed using this weighted similarity metric. For each cell, we consider the set

$knn_{r,i,1} \ldots knn_{r,i,200} \cup knn_{p,i,1} \ldots knn_{p,i,200}$ and identify the *k*-most similar cells within this set based on the weighted similarity metric as weighted nearest neighbors.

**Preprocessing details for each dataset**

*Cord blood mononuclear cells (CBMC) CITE-seq dataset:*

This CBMC dataset is a CITE-seq dataset from [23] and contains 8,617 cells with a panel of ten antibodies. We use the expression matrices as quantified in the original experiment. This experiment includes a small proportion of spiked-in murine 3T3 cells as negative controls. We apply SCTransform [15] to normalize gene expression data, and we apply a CLR transformation to normalize protein data within each cell. We use PCA to reduce the dimensionality of both datasets, taking 30 RNA and 7 protein dimensions to construct the WNN graph.

*Human bone marrow mononuclear cells (BMNC) CITE-seq dataset*

The BMNC dataset is a CITE-seq dataset from [16], consisting of 30,672 cells with a panel of 25 antibodies. We use the expression matrices as quantified in the original experiment. For gene expression, in order to facilitate comparisons with other methods, we use standard log-normalization with default parameters in Seurat. We apply a CLR transformation to normalize protein data within each cell. We use PCA to reduce the dimensionality of both datasets, taking 30 RNA and 18 protein dimensions to construct the WNN graph.

*ASAP-seq dataset of human PBMC*

We used the published human PBMC ASAP-seq dataset from [24], containing 4,725 cells with a panel of 227 antibodies, and performed processing with the Signac and Seurat packages. We use the ADT expression matrix, ATAC fragment files, and QC parameters from the original publication. We called peaks from the ATAC fragment files using the MACS2 callpeak function [25], and kept all peaks with -LOG10(qvalue) > 5 for the downstream ATAC analysis. We apply TFIDF to normalize ATAC peaks and CLR transformation to normalize protein data within each cell. We use LSI to reduce the dimensionality of ATAC normalized data, and PCA to reduce the dimensionality of protein. Then, we used LSI dimensions 2-50 LSI dimensions (excluding the first dimension as this is typically correlated with technical metrics in ATAC-seq data), and 30 protein PCA dimensions to construct the WNN graph.

*10x multiome ATAC+Gene expression dataset of human PBMC*

10x Genomics multiome (RNA + ATAC) data for human PBMCs was obtained from 10X website (https://support.10xgenomics.com/single-cell-multiome-atac-gex/datasets) and was processed using Signac and Seurat. ATAC-seq peaks were then identified for each cell type separately using MACS2, using the function CallPeaks in Signac 1.1.0 with arguments group.by='celltype' and additional.args='--max-gap 50'. Fragment counts for each peak were quantified per cell using the FeatureMatrix function in Signac. Per-cell quality control metrics were computed using the TSSEnrichment and NucleosomeSignal functions, and cells retained with a nucleosome signal score <2, TSS enrichment score >1, and total RNA counts <100,000 and >25,000. We apply SCTransform to normalize RNA counts and TFIDF to normalize ATAC peaks. We use LSI to reduce the dimensionality of ATAC data, and PCA to reduce the dimensionality of RNA. Then, we used 2-40 LSI dimensions and 1-40 protein PCA dimensions to construct the WNN graph. Motif analyses for the 10X RNA+ATAC and ASAP-seq datasets followed the suggested workflow described at https://satijalab.org/signac/articles/motif_vignette.html

*PBMC CITE-seq datasets of HIV Vaccine Trials Network samples*

Alignment and expression quantification: We applied standard pipelines to initially align and quantify the CITE-seq datasets newly generated for this manuscript. For both the 10x v3 (3' scRNAseq) and 10x Immune Profiling Solution (5' scRNA-seq), we used Cell Ranger 3.1.0 to align reads to the GRCh38 human genome with default settings. To quantify libraries of hashtag oligos (HTO) from cell hashing, or antibody-derived tags (ADT) from CITE-seq, we used Alevin [26]. A dictionary of barcode sequences for each antibody clone is included in Supplementary Table 1.

Quality control and doublet removal: We considered all cells that were detected in our RNA-seq, cell hashing, and ADT libraries. We first filtered out cells with that were outliers for the number of detected features from these modalities. We removed cells with < 500 detected genes, but also removed cells where we detected an aberrantly high number of features (more than 6,000 genes, more than 50,000 ADT reads, or more than 10,000 ADT reads), particularly to avoid clumps of antibodies that can occasionally attach to cells. We used our previously described hashing-based doublet detection strategy [5], implemented in HTODemux, to identify doublets that represent two or more cells representing different samples. Inspired by the scrublet package [27], we implemented a strategy to further remove doublets that may originate within the same sample (and would therefore not be identified through cell hashing). We first constructed a KNN graph based on the full ADT dataset, prior to removing cross-sample doublets. For each cell, we examined the percentage of neighbors that had been marked by HTODemux as doublets. If this percentage exceeded 20%, we reasoned that the cell's molecular profile was similar to a verified doublet, and therefore removed it from further analysis.

Sample integration (10X 3' CITE-seq experiments): To facilitate the identification of shared cell types across datasets, we applied our previously developed 'anchor' workflow [16] to integrate the datasets. We partitioned the dataset into 24 groups, each corresponding to one of the original samples representing one of eight volunteers, and one of three timepoints. To integrate the gene expression values, we first separately normalized each of the 24 groups using SCTransform, and applied the reciprocal PCA workflow, which is optimized for integration tasks with large numbers of samples and cells. When performing integration, we designated the unvaccinated cells (day 0), as reference datasets.

We integrated the protein measurements across samples using the same workflow, but after performing normalization within each cell using a CLR transformation. We did not perform variable selection on the protin data, but excluded four (out of 228) proteins from analysis: CD158, CD158b, CD158e1, and CD158f. We found that expression of these killer cell immunoglobulin receptors were highly variable across lymphocytes but did not correlate with other RNA or protein markers of cell state. As a result, they represented a 'nuisance' source of variation that diminished our ability to interpret clusters in the dataset. We therefore, chose to remove these proteins, analogous to the optional removal of cell-cycle stage-specific genes in scRNA-seq analysis, which can also represent nuisance sources of variation. We reduce the dimensionality of the integrated gene expression and integrated protein datasets via PCA. We use the top 40 and 50 dimensions respectively to construct KNN graphs from the RNA and protein modalities, which are used as input to the WNN procedure described above.

Clustering and annotation: To cluster our multimodal dataset, we first used the KNN graph based on the weighted RNA and protein similarities (referred to as the WNN graph), to calculate the Jaccard index (neighborhood overlap) between every pair of cells. This distance represents the edge weight in a shared nearest neighbor graph (SNN), which we used as input to the graph-based smart local moving (SLM) algorithm [28]. We initially clustered cells at a high resolution (resolution = 5), and performed differential expression (see below) on all pairs of clusters for both RNA and protein markers. We merged clusters that did not exhibit clear evidence of separation, or where the only differentially expressed features represented ribosomal genes or mitochondrial genes. In some cases (particularly for extremely rare cell types that required a higher resolution to be correctly annotated in our clustering), we increased the granularity of our clustering by subsetting cells in an individual cluster, and rerunning SLM on this subgraph. In our final annotations, we considered 57 total clusters.

We placed clusters into eight broad groups (Level 1 annotations: $CD4^+$ T cells, $CD8^+$ T cells, Unconventional T, B cells, Natural Killer (NK) cells, Monocytes, Dendritic Cells (DC), and Other (consisting of progenitors, circulating innate lymphoid cells, and additional rare populations expressing erythroid or platelet lineage markers). We further subdivided these groups into 30 Level 2 annotation categories representing well-described subtypes of human immune cells: $CD4^+$ T Naive, $CD4^+$ T Central Memory (TCM), $CD4^+$ T Effector Memory (TEM), $CD8^+$ TEM, etc., all thirty subtypes are listed at http://www.satijalab.org/azimuth). Our 57 clusters fall into subsets of these categories (i.e. $CD8^+$ TCM_1, $CD8^+$ TCM_2, etc.), and represent Level 3 annotations with the highest level of granularity (all listed in the legend for Figure 3C). We report markers for each of our Level 3 annotations in Supplementary Figure 5.

**Simulated addition of protein noise**

In Figure 2C, we perform a robustness analysis to explore the effects of artificially reducing the information content in one data type. To achieve this, we add increasing amounts of random noise to the protein data, immediately prior to running PCA. The amount of noise added is generated independently for each element in the matrix, and is drawn from a gaussian distribution with mean zero and increasing standard deviation (sd = 0.5, 1, 1.5, 2, 3, 4, 5). After adding noise, we repeated the WNN procedure.

**Comparisons with MOFA+ and totalVI**

In order to assess the performance of our WNN method alongside other recently proposed multimodal integration tools, we compared the results of WNN, Total Variational Inference (totalVI version 0.6.7) [29] and Multi-omics factor analysis v2 (MOFA+ version 1.1) [30], on the BMNC dataset. We followed the recommended settings and workflows for both methods, and further describe parameter choices below.

For totalVI, we use the RNA and ADT counts matrices as input. We use the subsample_genes function to select 4000 variable genes, and used 500 epochs for model training, as suggested in the totalVI tutorial (https://www.scvi-tools.org/en/stable/tutorials/totalvi.html). All other parameters were set to default settings. We identified nearest neighbors, and performed UMAP visualization on the learned latent space.

For MOFA+, we used the same normalization method as Seurat to facilitate direct comparison. As recommended in the MOFA+ tutorial (https://raw.githack.com/bioFAM/MOFA2_tutorials/master/R_tutorials/10x_scRNA_scATAC.html ), we used the z-scored data ('scaled' data) from the two assays as view1 and view2 for MOFA+. All other parameters were set to default or recommended settings. We identified nearest neighbors, and performed UMAP visualization based on the learned factors.

The UMAP plots in Supplementary Figure 2A-B show the results of all three methods (we also include independent RNA and protein analyses in Seurat for comparison). The plots show that the methods generally reveal similar sets of cell types, but with important differences. For example, regulatory T cells, defined by CD25 expression, are only separated in the WNN UMAP. Supplementary Figure 2B demonstrates that this is due to the fact that $CD25^+$ cells only form a distinct cluster in WNN analysis.

In order to move beyond visualization and quantify the performance of each method, we averaged the CD25 expression level for the calculated multimodal neighbors of each cell, returning a vector of predicted values. We quantified the performance of the method using the correlation (Pearson; Figure 2D, Spearman; Supplementary Figure 2), between predicted and measured values. For CD25, WNN analysis achieved the highest correlation, as cells that are $CD25^+$ are correctly identified as neighbors with other cells that are $CD25^+$ in the dataset. We repeated this analysis for all protein features, and found that, WNN analysis consistently achieved the highest correlation. We repeated the analysis for all transcriptomic features as well (Supplementary Figure 2), and observed similar performance for all methods. We note that transcriptomic correlations were also much lower, likely due to the substantial technical noise inherent to scRNA-seq data.

**TCR analysis**

To generate clonotype information for the 10X 5' samples, TCRαβ and TCRγδ fastq files were processed with cellranger vdj version 3.0.2 against the GRCh38 v2.0.0 reference as provided by 10x Genomics. Clonotype information was then manually added into Seurat as cell metadata, allowing us to explore the relationship between annotated cell type, molecular state, and TCR sequence.

**Identifying targeted immunophenotype panels**

For each of our 57 clusters, we aimed to identify a reduced set of antibodies that could enrich for cells in this molecular state. We utilized forward feature selection with balanced logistic regression to identify targeted surface protein markers for each cell type. This represents an iterative process where we successively add markers based on a greedy algorithm aiming to maximize the classification power of logistic regression. Prior to initializing the procedure, we randomly downsampled cells within abundant cell states to ensure that no cluster made up more than 5% of all cells in the dataset. We used the implementation for logistic regression with 5-fold cross validation in the boot R package [31]. We ran ten rounds of forward selection, allowing us to design panels of one to ten immunophenotypic markers for each cell type. To enhance the interpretability of these panels, we required the first five markers selected to have positive coefficients (i.e., to be enriched in the cell state of interest). These panels are reported in Supplementary Table 2. We used each panel to enrich for our 57 clusters 'in silico', using logistic regression with a decision boundary of 0.5 to set our gates. We report the enrichment, precision, and recall for each panel in Supplementary Table 2.

**Gradient analysis for NK and B cells**

In Figure 5G-H and Supplementary Figure 6, we identify genes whose expression level is correlated with a cell's position along a molecular gradient defined by a single protein. For example, in Figure 5G, we ordered cells along a gradient defined by CD16 protein expression. We then calculated Moran's I, a spatial autocorrelation metric proposed to identify trajectory-dependent genes in Monocle3 [32], to identify correlated genes. We plot a representative subset of these features in Figure 5G. We generate these plots by ordering cells on the x-axis based on their expression level for CD16 protein, and apply the ksmooth function from package stats with default bandwidth and parameters [33] to calculate smoothed gene expression levels across the trajectory. We utilize the same approach for trajectory analyses based on CD38 and CD27.

### Differential abundance of cell types across experimental conditions

In Figure 6 E-F, we analyze the composition of samples at different timepoints, and aim to find cell states whose abundance changes during the response. For Level 1 annotations, for each of the 24 samples, we calculated the percentage of each cell state in each sample, and ran two paired Wilcoxon tests: day 0 vs day 3, and day 0 vs day 7. No cell states exhibited significant changes. To search for more subtle changes, we calculated the relative abundance of all 30 Level 2 annotations in each sample within each Level 1 group (for example, for each sample we calculated the fraction of $CD14^+$ monocytes within the total pool of sample monocytes). These values were used as input to two paired Wilcoxon tests: day 0 vs day 3, and day 0 vs day 7. We detected significant shifts ($p < 0.05$) for three clusters, visualized for the 10X 3' samples in Figure 6F, with independent support for each of the signals in the 10X 5' datasets (Supplementary Figure 7).

### Identifying differentially expressed genes across cell types and experimental timepoints

In this manuscript (for example, Figure 4A), we identify differentially expressed (DE) genes and proteins that represent biomarkers of different cell states, or represent specific responses across experimental conditions. We used the wilcoxauc method from presto [34] to identify DE genes and proteins from our single-cell data. In addition, we performed the same DE test on the 'pseudobulk' expression values calculated from each of the 24 samples and aimed to identify markers that passed a stringent p-value threshold for both tests (adjusted p-value < $10^{-5}$). For space considerations, we typically report only the top 20 markers in each heatmap, and sort genes first by adjusted p-value and next by log fold-change to determine the top markers.

### Perturbation score

In Figure 6A, we aim to identify the cell types whose molecular state exhibits significant changes during the response to vaccination. We note that when calculating DE genes and proteins within a cell state, across experimental time points, the statistical power of these per-gene tests is heavily dependent on the abundance of the cell state. We therefore considered an alternative metric, the 'perturbation score' as described in [35], which quantifies the magnitude of the response across the transcriptome. To briefly summarize, we perform the following procedure to quantify the response for cells at day 3 vs. day 0 for each cell state. We first identify a set of genes that exhibit initial evidence of differential expression across timepoints but may not achieve statistical significance after multiple-testing correction (adjusted p-value < 0.1). We compute the pseudobulk expression of these genes after grouping cells by experimental timepoint, generating a vector representing the average expression of these genes for day 0 cells, and a second vector representing the average expression at day 3 cells. We define the 'perturbation vector' for this cell state as the difference between these two vectors, normalized to length 1. Finally, we project the transcriptome of each cell onto this vector and quantify the magnitude of this projection. We find that this approach helps to prioritize cell types that exhibit robust responses, particularly when comparing populations with vastly different abundances.

### Supervised principal component analysis for multimodal data

Due to the inherent levels of noise in single-cell RNA-seq, techniques such as PCA are often used to reduce the dimensionality of the dataset. PCA identifies correlated modules of genes, whose heterogeneous expression represent the largest sources of variance in the dataset. PCA is an unsupervised dimensional reduction technique, and while the correlated gene modules may typically represent markers of heterogeneous cell states in the dataset, they may also represent unwanted sources of variation related to technical noise, cell cycle state, or random fluctuations.

We therefore considered the application of supervised principal component analysis (sPCA) to our multimodal dataset. sPCA is a generalization of PCA that can be used not only for unsupervised learning, but also for regression and classification problems [36]. While PCA will identify the directions that explain maximal variance in the source data, sPCA can help pinpoint sources of variation that are of the greatest interest. To accomplish this, sPCA takes as input a kernel which describes the similarity between any two cells based on a response outcome. We set this kernel to represent the Jaccard distances derived from our WNN graph, as this considers the response outcome to be the weighted combination of RNA and protein profiles. sPCA will then estimate a set of principal components that have maximal dependence on the response variable [36]. These components should represent the optimal transcriptomic vectors that can be used to separate the cell types defined in our multimodal dataset. Therefore, the sPCA procedure can identify the set of principal components that can transform the data in a single modality to best capture the structure in a multimodal dataset. We emphasize that sPCA takes as input a cell-cell similarity kernel, but does not require cells to be labeled or placed into discrete clusters. Therefore, it can capture both discrete and continuous sources of variation in a multimodal dataset.

Formally, sPCA transforms the dataset to maximize the dependency with the response variable. We implement the method described in [36], where the Hilbert-Schmidt Independence Criterion (HSIC) is used as the dependency measure. To apply this method in the context of single cell multimodal data, we define the following:

$X$: data matrix for gene expression measurements
$Y$: data matrix for protein measurements.
$U$: Transformation of $X$ (for example, a set of principal components)
$K$: Kernel derived from $U$, describes the cell-cell similarity in $X$
$L$: Kernel derived from the WNN graph, and describes the cell-cell similarity based on a weighted combination of $X$ and $Y$

The HSIC between two kernels $K$ and $L$ is:

$$HSIC(K, L) = \frac{1}{(n-1)^2} tr(KHLH)$$

The goal of the sPCA is to identify $U$ that maximizes $HSIC(K, L)$

$$HSIC((U^T X)^T U^T X, L) = \frac{1}{(n-1)^2} tr(X^T U U^T XHLH)$$
$$= \frac{1}{(n-1)^2} tr(U^T XHLHX^T U)$$

As described in described in [36], the optimization problem reduces to:

$$\underset{U}{\text{argmax}} \quad tr(U^T XHLHX^T U)$$
$$subject\ to\ U^T U = 1$$

where H is the centering matrix $H_{ij} = I - n^{-1}ee^T$.

This optimization problem has a closed form solution. $U$ represents the eigenvectors of matrix $XHLHX^T$, based on the top $d$ eigenvalues, where $d$ represents the desired number of components. Each vector in $U$ describes the relative importance for each gene in defining this component (i.e, $U$ represents a set of feature component loadings).

**Mapping query datasets to a multimodal reference**

We compute the sPCA transformation described above for our reference dataset, and can subsequently project this transformation onto any query dataset consisting of PBMC. This enables us to perform supervised analysis of the query datasets. Since our sPCA was computed based on a reference defined by a large number of cells and antibodies, this transformation will likely be more informative than an unsupervised PCA computed on a new scRNA-seq query. This transformation should therefore be more capable of separating cell types in the query dataset. As a secondary benefit, projecting the sPCA transformation onto a query dataset places the query in the same low-dimensional space as the reference. This provides a starting point to integrate the two datasets, which can assist in the visualization and annotation of the query as described below.

**Reference-based Integration for query datasets**

In [16], we demonstrate a workflow to identify reference-based transfer 'anchors' between reference and query datasets. Briefly, this workflow first projects a transformation calculated on the reference dataset onto the query. The method next identifies mutual nearest neighbors [37] between the reference and query datasets, based on this L2-normalized low-dimensional space. These anchors can be used to transfer discrete or continuous data from the reference onto the query. For cell annotation (transfer of discrete label from reference to query), each query cell is assigned a label based on a weighted vote classifier, where each anchor provides a vote that is weighted by its similarity to the query cell. When classifying cells in this manuscript, we apply the same workflow, but use the sPCA transformation described above for projection.

In [16], we also provide methods to leverage an existing set of anchors in order to modify the underlying gene-expression levels, allowing shared cell types to cluster together across experiments. We apply a similar workflow here. However, instead of correcting values in high-dimensional space, we correct in low-dimensional space. This substantially improves the speed of the method. Moreover, at the conclusion of this procedure, we have placed the query dataset in the same low-dimensional representation (defined by the sPCA transformation) as the reference.

Having placed both the query and reference dataset in the same space, we have two options to visualize the query dataset. The first is that we can project the query data onto the same UMAP projection as has been previously computed. To accomplish this, we use the umap_transform functionality implemented in the R uwot package, which enables new points to be adding to an existing embedding. We use this approach to project query datasets onto the reference-defined visualization shown in Figure 3D. Together, these methods enable a fully automated pipeline to leverage a multimodal single-cell reference to annotate and visualize new single-cell query datasets, even if only the transcriptome was measured. To facilitate users applying this approach to interpret their own datasets from either healthy or diseased PBMC, we have provided a web application (http://www.satijalab.org/azimuth) to automate these analyses.

A second option for visualization is to compute a new ('de-novo') UMAP visualization, which can be computed after merging the reference and query datasets together. For the analysis of the COVID-19 dataset [38], we compute both visualizations. The reference-based UMAP is shown in Figure 7D, while the de-novo UMAP is shown in Supplementary Figure 8. One advantage of the de-novo approach is that it can help to visualize populations in the query that cannot be effectively represented in the reference. For example, the Wilk et al. dataset contains subsets of developing and differentiated that were not captured in our reference, as well as subpopulations of monocytes whose expression profiles are heavily perturbed in COVID-19 samples. In the reference-based visualization, the umap_transform functions aims to embed these cells adjacent to their closest neighbors in the reference, which often places these cells at the boundary of cell clusters. In the de-novo visualization, all three of these populations remain distinct from reference cells even after integration (Supplementary Figure 8). We encourage users to compute both visualizations to understand how their dataset can be interpreted in light of a reference, and also to flag any particular populations that may not be well represented.

We leverage this reference-based mapping workflow to interpret the 5' scRNA-seq datasets generated for this manuscript. We use the same QC, normalization, and doublet filtration procedures to analyze the 5' data as described earlier in this section. We apply the reference-based integrative analysis procedures described above to project the 5' scRNA-seq data onto the UMAP visualization defined by the 3' dataset, and also to transfer discrete labels. The annotation and projected UMAP are shown in Supplementary Figure 6F, while the UMAP visualization with annotated clonotype structures is shown in Figure 5K.

Similarly, we applied the same pipeline to map the CITE-seq datasets from [39]. We downloaded the dataset at https://doi.org/10.35092/yhjc.c.4753772, applied SCTransform normalization, and repeated the mapping procedure applied above. While the dataset contains measurements for 82 proteins alongside the transcriptome, we used only the transcriptome for reference mapping and the transfer of Level 2 annotations. This allowed us to use the withheld protein data for benchmarking with scArches (version 0.1.2) [40], as shown in Supplementary Figure 8.

**Benchmarking Seurat reference-mapping with scArches**

To run scArches, we followed the tutorial released by the authors. We first integrated our 24 3' scRNA-seq samples into a reference atlas, using the same variable genes as used in the WNN analysis. We obtained poor results with the default nb loss function, and as suggested in the tutorial, tried the sse loss function as an alternative. We trained the scArches model using recommended parameter settings of 150 epochs and a batch size of 128, and next mapped query cells onto the reference using recommended parameters in the tutorial. To facilitate fair comparisons between our reference mapping workflow and scArches, we forced both methods to return the most likely annotation for each query cell.

We note the extensive challenges in benchmarking reference-based annotation workflows in the absence of ground-truth cell labels. By withholding the protein data from consideration during the mapping process, we can use the protein measurements as an independent assessment of prediction quality. For 35,619 cells (67.1%), Seurat and scArches returned the same annotation. For the remaining 17,480 query cells, the two methods returned two divergent annotations (for example, suppose that Seurat annotated the cell as CD4 Treg, and scArches annotated as NK). In the reference dataset, we calculated the protein centroids for the CD4 Treg and NK clusters. We then calculated the Pearson correlation between these centroids, and the protein values for the individual cell. If the cell's protein levels exhibit a high correlation with the centroid of CD4 Treg, but a low correlation with the centroid of NK, this suggests that the Treg annotation is correct. This metric and approach are inspired by scmap [41]. Essentially, in cases where two methods disagree based on an RNA classification, we attempt to classify the cell based on its protein levels to see if there is strong evidence for one annotation vs another. In 73.8% of cases, we observe stronger support for the Seurat annotation (Supplementary Figure 8).
